# Complex ecosystems lose stability when resource consumption is out of niche

**DOI:** 10.1101/2023.11.30.569477

**Authors:** Yizhou Liu, Jiliang Hu, Hyunseok Lee, Jeff Gore

**Affiliations:** Physics of Living Systems, Department of Physics, MIT, Cambridge, MA 02139, USA; Department of Mechanical Engineering, MIT, Cambridge, MA 02139, USA

**Keywords:** ecology, resource, stability, dynamics

## Abstract

Natural communities exhibit diverse dynamics, encompassing global stability, multi-stability, periodic oscillations, and chaotic fluctuations in species abundances. Resource-consumer interactions provide a lens for mechanistic understanding of community behaviors, yet only a globally stable equilibrium can exist in the original MacArthur resource-consumer model. Here we find that diverse dynamics emerges when species consume resources that contribute little to their own growth. Key to understanding these results is comparison of niche range (difference between growth-promoting resources of similar species) and consumption range (difference between growth-promoting resources and the resources that are actually consumed). If the consumption range is small, we observe global stability as in the MacArthur model. But when consumption range increases to about the niche range, stability is lost, giving rise to emergent alternative stable states, globally stable states but with species extinction, and slightly later, persistent fluctuations. Given its importance in dictating stability, we define the ratio between consumption and niche ranges as encroachment, and find that it predicts key community properties like diversity and attractor basins even after the instability transition. In particular, after the loss of stability (encroachment greater than one), species extinction increases approximately linearly with encroachment. Since encroachment relies only on intrinsic species properties, community stability is resilient to environmental changes such as resource supply and mortality rate. Encroachment provides a framework for capturing resource competition due to growth-consumption inconsistency in complex communities as well as a robust quantitative characterization of the resulting emergent dynamics.

## Introduction

Ecological systems exhibit a range of dynamical behaviors which are important to community function. From gut microbiota to grass land plants, constant abundances over decades are signatures of global stability [1–3]. Some natural communities, however, may display alternative stable states [4–9], leading to the possibility of becoming “stuck” in undesirable states that may be associated with diseases [5]. Communities have also been observed to display persistent fluctuations in species abundance, including limit cycles [10–12] and chaos [13–15]. Despite the importance of these community dynamics, we have limited understanding of the factors that determine how a community behaves.

Given the complex nature of ecological systems, phenomenological descriptions (general Lotka-Volterra model) supply basic insights into different dynamical behaviors. Pioneering works probing the stability of an equilibrium have concluded that increasing species number and interaction strength will typically destabilize a community [16,17]. Recent theory predicts how key features of an assembled community (e.g., expected diversity) depend on interspecies interaction strengths [18,19]. Combined with the understanding of stability, a complete picture encompassing full coexistence of species, partial coexistence, and instabilities has been established and experimentally verified [20]. Beyond the transition to instability, researchers are working to capture alternative stable states [21,22] and various fluctuating behaviors [23–26] in more detail. Despite these successes in understanding, coarse-grained pictures built on abstract strengths of interaction and types of interactions (e.g., cooperative, competitive, and neutral) are limited in linking biological mechanisms and community behaviors, restricting our ability to predict and control ecosystems.

Resource competition is a key interaction in many natural communities [27–29]. A better understanding of how species and resources alter community dynamics would enable directed interventions such as therapies to restore gut microbiota to desired states [5]. The simplest resource explicit models only include resources and consumer species. Species consume different resources to grow, and competitive interactions emerge since resource consumption hinders the growth of other species. Historically, efforts on resource-consumer models have focused on the important question of feasibility [30–33], i.e., how to obtain high-diversity fixed points with non-negative abundances and concentrations, motivated by the disagreement between the competitive extinction principle [34,35] and the reality of high diversity under few distinct resources (the “paradox of the plankton”). Trade-offs in metabolism due to fixed enzyme budgets can beat the competitive extinction principle [30]; noises in resource supply and diversity of species will change the dependence of equilibrium resource concentrations on external supplies [31]; and different ways of supplying resources affect expected diversities of feasible fixed points [32] …. After obtaining such a feasible coexistence equilibrium, we often only see global stability displayed as the original MacArthur resource-consumer model [36]. Understanding feasibility alone cannot lead to explanations of various dynamics exhibited in natural systems. Learning from the development of phenomenological results, instability is the key to rich dynamics. In the context of resource explicit models, instability can be induced by cross-feeding [37], external noise [38], and increasing non-linearity [38,39], yet the mechanistic essence behind instability transitions driven by resource competition has not been well characterized.

In this study, we explored a generalization that allows for variations in yields (biomass growth over resource consumed) among species and among resources [40]. The global stability in the MacArthur model relies on the implicit assumption that biomass yields are essentially determined by resources, i.e., the efficiency that each resource is converted to biomass is constant across species. Once we allow variations in yields, the consequences are well known for simple communities with two species and two resources, where bistability rather than global stability can occur when each species consumes a great deal of the resource that the other species grows well on [41]. However, the result of heterogeneity in yields is not well understood for large complex communities containing many species and many resources.

Here we find that the full spectrum of dynamics (e.g., chaos, limit cycles, and alternative stable states) emerge under sufficiently large variation in yields which leads to a deviation between the resources that a species consumes and those that are most important for growth. This deviation can be quantified by comparing niche range (difference between growth-promoting resources of similar species) and consumption range (difference between growth-promoting resources and the resources that are mainly consumed). We find that the degree of stability is determined by the ratio of consumption range to niche range, defined as encroachment. The communities are globally stable when encroachment is small. Communities become unstable when encroachment surpasses 1, and we observe various dynamical behaviors including alternative stable states, global stability after species extinction, and persistent fluctuations. Beyond stability, we find encroachment can further predict key community properties like diversity loss and attractor basins. The seemingly scattered yet rich dynamical behaviors driven by resource competition can be organized along a single dimension, the encroachment. Finally, we discuss how encroachment can help to unify previous results under one mechanistic picture.

## Results

We employed the simplest resource-competition framework to investigate interactions between species and resources. Time evolution of resource concentrations is controlled by how species consume resources and how resources are supplied (Fig. 1A):

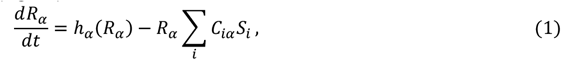

where *R*_*α*_ represents concentration of the *α*th resource, *S*_*i*_ denotes abundance of the *i*th species, *C*_*iα*_ is the per capital consumption rate of species *i* for resource *α*, and *h*_*α*_(*R*_*α*_) is a function encoding resource supply. Species dynamics rely on how species grow based on resources and die according to mortality rates (Fig. 1B), as described by

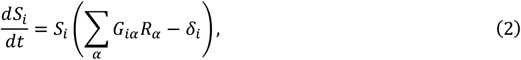

where *G*_*iα*_ is the per capital growth rate of species *i* with unit resource *α* and *δ*_*i*_ the mortality rate of species *i*. The matrices *G* and *C* will be referred to as growth rates and consumption rates, respectively. Biomass yields are rigorously defined as *G*_*iα*_/*C*_*iα*_, encoding how efficiently each resource will be converted to biomass. Such linear uptake models trace back to MacArthur’s work [36] in the context of zoology, where *h*_*α*_(*R*_*α*_)=*g*_*α*_*R*_*α*_(*K*_*α*_ − *R*_*α*_) takes the form of logistic growth for biotic resources like grass. For microbial communities, chemostat resources are commonly considered, taking the form *h*_*α*_(*R*_*α*_)=*l*_*α*_(*κ*_*α*_ − *R*_*α*_) where *κ*_*α*_ can be understood as concentration at the source and *l*_*α*_ as the dilution rate. In the main text, we present results of simulating constant resource supply, *h*_*α*_(*R*_*α*_)=*γ*_*α*_, and leave those of other resource supply methods to supplementary information (SI) [42]. Interestingly, the instability transition is similar for the different methods of modeling resource supply (Fig. S1).

**Fig. 1.**
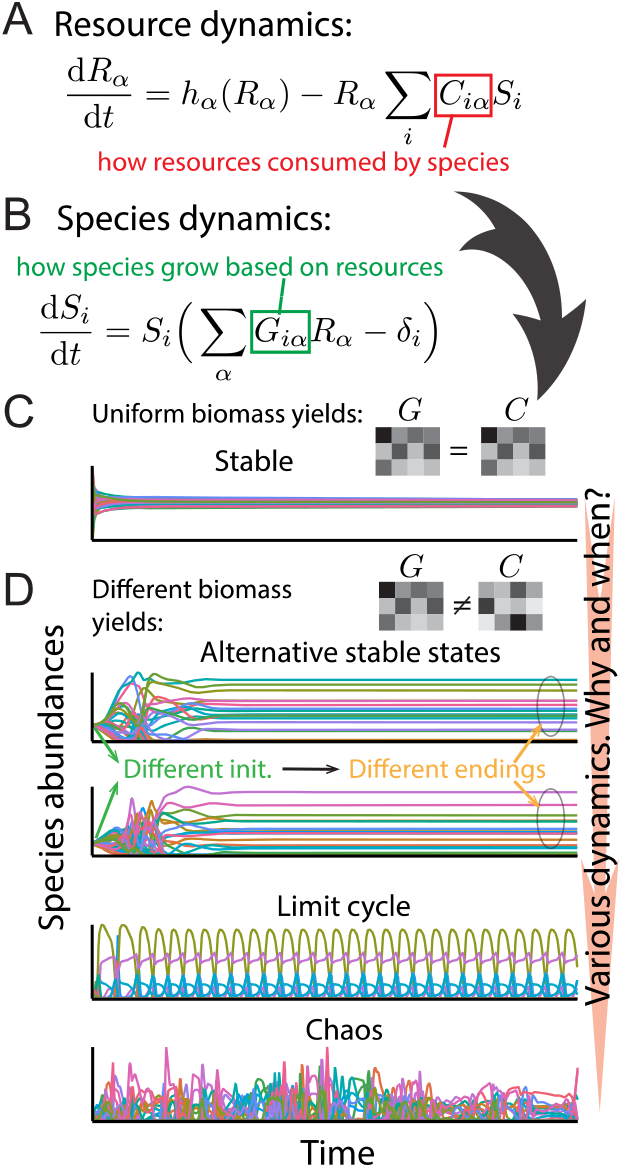
Divergence between consumption and growth can lead to instabilities and subsequently rich dynamics. (A) Dynamics of resource concentrations are governed by how species consume resources, encoded in the matrix *C*, and how resources are supplied. (B) Species abundance changes are subject to how species grow based on different resources, as encoded in the matrix *G*, and the mortality rates. (C and D) Various possible dynamics can happen when we change the correlation between consumption rates *C* and growth rates *G*. Given a feasible fixed point, it is stable when *C*=*G* as shown in (C). When *G* ≠ *C* (displayed in D), alternative stable states can appear in the same community, and it is possible to observe persistent fluctuations including both limit cycles and chaos. Detailed simulation settings are in SI [42].

We began by characterizing the range of dynamical behavior displayed by communities interacting via this simple mode of resource competition. We considered a community with *N*_*S*_ species and *N*_*R*_ resources with an interior fixed point (that may or may not be stable). As expected from previous work [36,37], when yields are constant (*C*=*G*) the fixed point is globally stable and all communities reach the same final state independent of the initial concentration of each of the species and resources (Fig 1C). As the correlation between growth rates *G*_*iα*_ and consumption rates *C*_*iα*_ decreases we begin to observe alternative stable states, as well as non-converging behaviors like limit cycle oscillations and chaotic fluctuations (Fig. 1D). This simple model therefore displays the full range of dynamical behaviors that have been observed in natural communities, including global stability, alternative stable states, limit cycle oscillations, and chaos. Although it is evident that divergence between *G* and *C* can lead to instabilities and subsequently various dynamics, it is unclear what is the essential driver behind these transitions.

To gain insights into the origin of the different dynamical regimes of these communities, we started by considering the local stability of a given fixed point as has been done previously for phenomenological models [16,17]. In the study of local stability, we will introduce a special situation where resource concentrations are equal at the fixed point. This special case can help to build basic geometric intuitions for understanding stability. Later, we will consider randomly sampled fixed points and generalize the geometric understanding to reveal the topological essence of local stability.

We started the exploration of instability by looking at special fixed points with equal resource concentrations. The equal concentrations can be understood as a consequence of metabolic trade-offs [30]. The simplest metabolic trade-off is given by

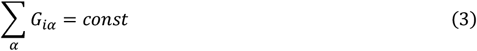

for all *i*. This constraint on of *G* could arise if different species have the same fixed enzyme budget, meaning that if one species allocates more enzymes for certain resources then it will grow more slowly on other resources. The mortality rates in this case are the same, which can be interpreted as a universal dilution rate in a chemostat. Combining the assumptions, a feasible fixed point of resources *R*^*\**^ can always be ensured. It can be readily verified that different resources have equal equilibrium concentrations, i.e., 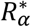 are the same for different *α*.

We first sought intuition by considering a simple community with two species and two resources [41]. Here, growth rate vectors and impact vectors determine local stability. A growth vector *G*_*i*_, is the *i*th row of the matrix *G*, denoting the resources species *i* grows well on. Impact vector of species *i, V*_*i*_, defined by 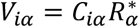 (per capital consumption rates at the fixed point), as its name indicates, describes how species *i* impacts resources via consumption. In the resource space, the system is regarded to be most stable when growth vectors are aligned with the corresponding impact vectors. This is because growth vector *G*_*i*_ is the normal vector of the zero net growth isocline (ZNGI) of species *i*, and *V*_*i*_ ∥ *G*_*i*_ means the species tries to drag resources back to equilibrium by consumption in the rapidest direction (Fig. S2 explains details of Tilman’s graph [42]). With uniform equilibrium resource concentrations, 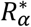 are the same, making *V*_*i*_ parallel with *C*_*i*_, the *i*th row of the matrix *C*. The case *G*=*C* then leads to *V*_*i*_ ∥ *G*_*i*_ for all *i*, ensuring stability. However, if species consume more of the resource that they grow poorly on, then the fixed point can become unstable, leading to bistability in the simple two-species community [41]. In these graphic representations, the instability transition is vividly captured by a flipping of impact vectors.

Unfortunately, Tilman’s graphical approach has no direct generalization to complex communities, but we followed the key intuition to conjecture one. The first fact is that only directions of the growth and impact vectors matter (see Tilman’s graph in SI [42]). Therefore, we can normalize the vectors to the simplex (a vector *x* is on the simplex if *∑*_*α*_ *x*_*α*=_1). For example, a community growing on three resources has a simplex that is a triangle that cuts through three-dimensional resource space (Fig 2A). The normalized growth vector is given by 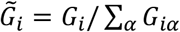 which keeps the direction but eliminates magnitude information (Fig. 2, B and C). Normalized impact vector 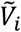 is obtained in the same way. The second key insight from Tilman’s graphs is that stability depends on whether impact vectors are more aligned with their own growth vectors or not. To quantify the degree of alignment, we define consumption ranges using the distances between normalized impact vectors 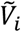 and the corresponding growth vectors 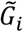 (radius of dashed circles in Fig. 2B). The critical degree of alignment is captured by the distances between neighboring normalized growth vectors (line between squares in Fig. 2B). For species *i*, we define the half distance between 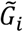 and 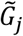 which belongs to its nearest neighbor species *j* as the niche range of species *i* (niche range uses the length distance to evaluate the similarity between niches, and previous works on niche overlap used inner product between the vectors for the same purpose [43,44]). Once the consumption range is greater than the niche range, flipping of impact vectors is enabled to occur, i.e., one’s consumption vector is more aligned with another one’s growth vector while the other one’s consumption vector is closer to one’s own growth vector (e.g., Fig. 2C). The ratios of consumption ranges and niche ranges turn out to be essential, which we will call “encroachment” as it quantifies the degree to which species consume resources that promote the growth of other species. To identify community level collective behaviors, we define the simple encroachment *E*_*s*_(*G, V*) based on the geometry as the averaged ratio of consumption range to niche range over species:

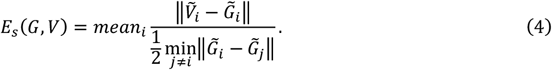

Note that *E*_*s*_(*G, V*) > 1 is only a necessary condition for instability since it enables flipping rather than ensures flipping. In high dimensional cases, there are more configurations leading to instabilities besides pairwise flipping, which, however, still may be able to occur only if *E*_*s*_(*G, V*) > 1 (Fig. S3 in SI [42]). We have proposed a simplified condition for local instability inspired by Tilman’s graph, quantifying how species in a community encroach on each others’ niches.

**Fig. 2.**
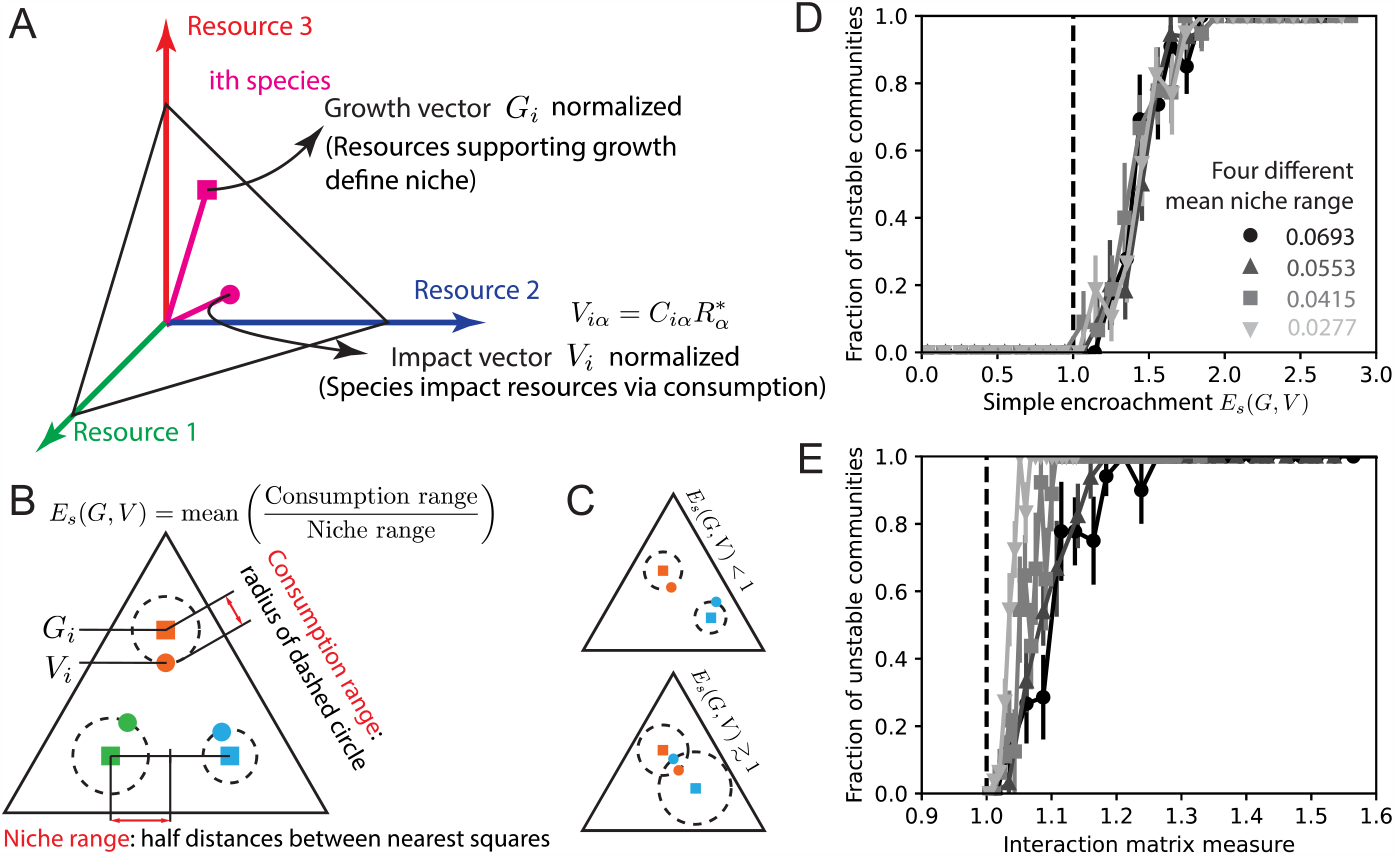
Niche range and consumption range defined based on directions of species growth and resource consumption well capture instability transition for communities with equal equilibrium resource concentrations. (A) We use a three-resource example to explain graphic notations. For the *i*th species, the growth vector *G*_*i*_ and impact vector *V*_*i*_ are three dimensional vectors with positive elements. After normalization to the simplex which is the triangle in (A), we use a square to denote the end of the growth vector, and a circle for the impact vector. (B) On the simplex, the normalized growth vectors represent niches. We use minimum half distances between normalized growth vectors as a measure of the range of niche. And the distance between one’s normalized growth vector and impact vector is used to define consumption range. At last, the mean ratio of consumption range to niche range over species is defined as simple encroachment for the community. (C) Illustrated examples with two species and three resources, where the upper one has encroachment *E*_*s*_(*G, V*) < 1 (i.e., consumption range is smaller than niche range, and the community is stable), and the lower one has *E*_*s*_(*G, V*) > 1 (i.e., consumption range surpasses niche range, and the community is unstable). (D) Numeric results for four classes of communities with different mean niche ranges support the validity of *E*_*s*_(*G, V*) in identifying instability. (E) Criterion from the simplest random matrix analysis fails to identify the instability transition. The interaction matrix measure is obtained via random matrix analysis of a reduced local interaction matrix based on the local mapping from the resource-consumer model to generalized Lotka-Volterra dynamics [42]. Numeric tests are done for communities with *N*_*S*=_16, *N*_*R*=_32 [42], and stability in (D) and (E) is rigorously inferred from the local Jacobian. Error bars represent SEM.

To test whether simple encroachment *E*_*s*_(*G, V*) can indeed predict the stability of complex communities, we randomly sampled communities with uniform equilibrium resource concentrations and determined their stability. The growth rates *G* were uniformly sampled on different regions of the simplex. Consumption rates *C* were perturbed from *G*, and species abundances at the given fixed point are uniformly sampled from [0,1] (detailed sampling and simulating schedule are in SI [42]). Since we can sample *G* from larger or smaller regions of the simplex, we can tune the magnitude of niche ranges. We show representative results (Fig. 2D) obtained from fixed points with 16 species, *N*_*S*=_16, and 32 resources, *N*_*R*_ *=* 32 (*N*_*S*_ /*N*_*R*_=1/*2* is the maximum for randomly assembled communities in chemostats [32]), and will discuss the high-order effects of *N*_*S*_ /*N*_*R*_. Consistent with our prediction that encroachment reflects the stability of complex communities, our numerical stability analysis indicates that communities lose stability near *E*_*s*_(*G, V*)=1 (Fig. 2D). The degree of stability is determined by the Jacobian at the given fixed point. Regardless of how different the growth vectors are (value of mean niche range), as long as *E*_*s*_(*G, V*) is the same, the fraction of unstable communities is almost the same. Since *E*_*s*_(*G, V*) > 1 is a necessary condition, there can be stable communities after it surpasses 1. We therefore find that the simple encroachment *E*_*s*_(*G, V*) accurately predicts the stability of complex multispecies communities with equal equilibrium resource concentrations.

Resource-consumer models can be mapped to a local general Lotka-Volterra type model near the fixed point [42], enabling alternative ways to identify stability. With the help of this bridge, we can get a phenomenological interaction matrix based on growth rates, consumption rates, and the species abundances and resource concentrations at the fixed point. Stability analysis of such interaction matrices using random matrix theory have been well established [16,17,45]. We find that the stability criterion derived from random matrix theory fails to predict the stability of our communities. The criterion predicts that the instability transition will occur sharply when the interaction matrix measure is 1 (definition and derivation in SI [42]), but all the stable communities have interaction matrix measure greater than 1 (Fig. 2E). Moreover, for different mean niche ranges, the instability transitions occur at different values of the interaction matrix metric, indicating that this measure alone cannot capture the instability transition. This failure arises because random matrix predictions assume correlations in the interaction matrix are simple (lacking higher-order correlations), while this is not true for resource-consumer models even when the correlations between *G* and *C* are simple (e.g., each *G*_*iα*_ correlates with *C*_*iα*_ and the correlation is constant for all *i, α*). In summary, although the simple encroachment *E*_*s*_(*G, V*) is highly simplified and can only yield a necessary condition for instability, in practice it captures the transition to instability better than estimates derived from random matrix theory.

After the success in the realm with equal equilibrium resource concentrations, we moved to explore encroachment and stability criterion for general fixed points. Without uniform equilibrium resource concentrations, the fact that the impact vectors depend on resource concentrations 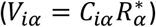 can lead to large differences between the directions of intrinsic species consumption rate, *C*_*i*_, and impact vector at the fixed point, *V*_*i*_. Under such a situation, *E*_*s*_(*G, V*) can be larger than 1 even when *G*=*C*, leading to a failure of predicting stability. This effect can be observed by simulating communities at four fixed-point resource concentrations with different evenness (Fig 3A). As the equilibrium concentrations become increasingly non-uniform the instability transition is delayed, i.e., *E*_*s*_(*G, V*) needs to be larger than a threshold value greater than 1 (e.g., squares in Fig. 3A), and eventually *E*_*s*_(*G, V*) totally loses the ability to capture instability (circles in Fig. 3A). Non-uniform equilibrium resource concentrations can lead to the failure of *E*_*s*_(*G, V*) to predict stability by separating the impact vectors and growth vectors, not changing whether the community is stable not or not yet increasing consumption range. Our definition of simple encroachment therefore fails to predict the instability transition for arbitrary fixed points.

**Fig. 3.**
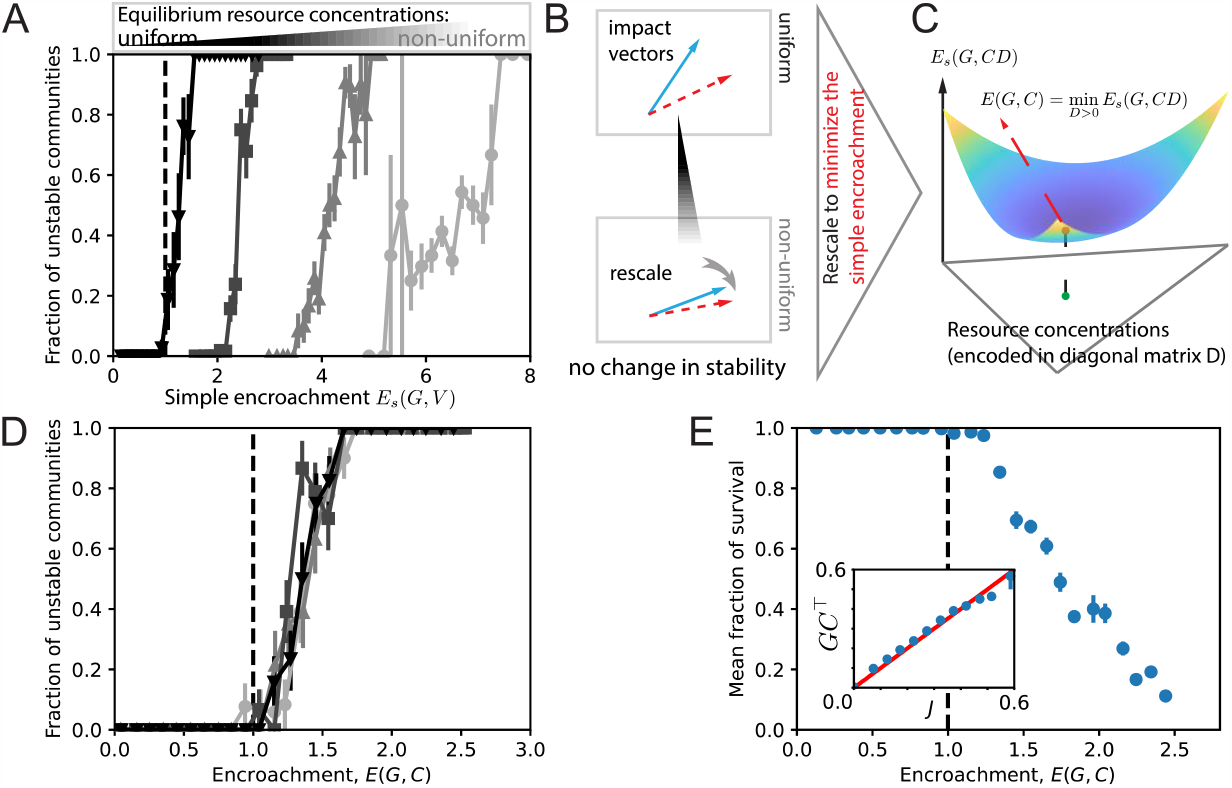
Stability is robustly irrelevant to where the interior fixed point is, leading to refined encroachment for general cases and new stability criteria. (A) We test four classes of communities, where within each class, communities have the same equilibrium resource concentrations. The measure *E*_*s*_(*G, V*) fails to predict stability correctly with non-uniform equilibrium resource distributions. (B) Non-uniform equilibrium resource concentrations do not change relative relationships among impact vectors, indicating that stability is irrelevant to fixed-point position but intrinsic growth rates *C* and growth rates *G*. We therefore conclude that if the community can be stable at certain point based on previous geometric arguments (not necessarily the real fixed point), topologies of growth rates *C* and growth rates *G* are similar enough and local stability can be ensured at the real fixed point. (C) We use optimization techniques to find the concentration distribution where system is most likely to be stable, which is named as the virtual fixed point, and define encroachment as the simple encroachment evaluated there. (D) The newly refined encroachment, *E*(*G, C*) works well on the same numeric data in (B), supporting the topological essence of stability. (E) Unlike (B) and (D) where stability of communities is determined by Jacobian, we simulate the dynamics starting near the fixed point to test stability. Fraction of survival species as evidence supports the stability criterion based on *E*(*G, C*). Moreover, the value of *E*(*G, C*) can predict that of mean survival fraction. The inner small figure shows the mean fraction of unstable modes (number of unstable eigenmodes over that of all eigenmodes) of *GC*^*T*^, reduced from local Jacobian inspired by the fact stability is largely irrelevant to local information, is almost the same as that of the local Jacobian *J*. Most numeric results are obtained when *N*_*S*=_16, *N*_*R*=_32, while for *GC*^*T*^, we tested different *N*_*S*_ [42]. Error bars represent SEM.

Revisiting the simple communities inspired a possible solution by defining a modified measure for stability. Rewriting impact vectors 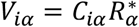 in the matrix form, we have *V*=*CD*(*R*^*\**^), where *D*(*R*^*\**^) is a diagonal matrix filled by the vector of equilibrium resource concentration, *R*^*\**^. In the two-resource cases, the transformation *V*=*CD*(*R*^*\**^) simply rescale each *C*_*i*_ vector towards the same direction, but will not change the relative relations among different *C*_*i*_. That is, whether impact vectors flip (with respect to the growth vectors) or not is irrelevant to *D*(*R*^*\**^) and is determined by whether some relative topology among *C*_*i*_ is similar enough to that of *G*_*i*_. For general complex systems, we assume this argument that stability is irrelevant to where the interior fixed point is, to obtain a trick of virtual fixed point and quantify how close the relative relationships among *C*_*i*_ are to those of *G*_*i*_. Let *D* be any positive definite diagonal matrix whose elements are interpreted as resource concentration of a fake fixed point, we can take *CD* as the impact vector at the fake fixed point and calculate *E*_*s*_(*G, CD*). If minimum *E*_*s*_(*G, CD*) over possible *D* can be smaller than 1, it means there exists a resource concentration at which the community is locally stable based on the previous geometric understanding. With the assumption that local stability is irrelevant to where the fixed point is, we can conclude that the system is stable at its real fixed point *R*^*\**^. On the contrary, if the minimum *E*_*s*_(*G, CD*) is greater than 1, then the system cannot be stable anywhere with those *N*_*S*_ species and *N*_*R*_ resources, and it is certainly not stable at the real fixed point *R*^*\**^. With this trick of virtual fixed point, we realize the minimum value of *E*_*s*_(*G, CD*) may be a general indicator of stability which quantifies the similarity in topologies of the growth vectors and consumption vectors. By refining encroachment as

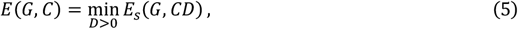

we can state our stability criterion as the community is locally stable if *E*(*G, C*) < 1. In a three-resource space, Fig. 3C illustrates the definition of *E*(*G, C*) and conceptually where the real and virtual fixed points are (detailed optimization processes are explained in [42]). Using the same set of communities (in Fig. 3A) to test *E*(*G, C*), we found instability transition is well captured regardless of the evenness of equilibrium resource concentrations (Fig. 3D). Our refined metric of encroachment *E*(*G, C*) is therefore able to accurately predict instability of complex communities without knowing the specific fixed-point position.

The fact that stability does not depend on fixed-point position is not obvious from canonical Jacobian analysis, which inspired us to eliminate the dependence of local fixed point in the Jacobian and obtained a characteristic matrix *GC*^*T*^ [42]. Defining unstable mode fraction of a Jacobian as the ratio of the number of its unstable eigendirections to the number of species, *N*_*S*_, we observe an elegant result that unstable mode fraction of *GC*^*T*^ is the same as that of the real Jacobian on average (internal subplot of Fig. 3E). We therefore find, perhaps surprisingly, that stability only depends on *GC*^*T*^. If we further assume *G* elements to be i.i.d., *C* elements to be i.i.d, and *G*_*iα*_ and the corresponding *C*_*iα*_ have the same correlation *Y*, random matrix theory for this type of *GC*^*T*^ matrix [46] provides a concise stability criterion:

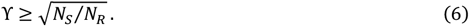

The random matrix result is obtained in the limit *N*_*S*_ and *N*_*R*_ go to infinity while the ratio *N*_*S*_*/N*_*R*_ is kept constant. The correlation *Y* uses inner product to quantify similarity between one’s consumption and growth vectors (corresponding to consumption range), and 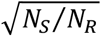 is an estimated bound of niche overlap (corresponding to niche range). A greater “inner product” *Y* refers to a smaller consumption range (distance between vector ends), and thus leads to stability. The operation of eliminating means in calculating *Y* suffices for the assumed *G*_*i*_ and *C*_*i*_ in Eq. (6) to be aligned, which is like *C → CD* in calculating *E*(*G, C*). To conclude, stability is found to be related to topologies of growth and consumption vectors, and following such intuition we can obtain one more characteristic indicator for stability, *GC*^*T*^, simplifying the original Jacobian analysis.

We next sought to determine how much species extinction occurs after communities lose stability (encroachment *E*(*G, C*) > 1). Unlike previous numeric tests which only evaluated the local Jacobian to infer stability, here we examine stability by simulating the dynamics (Eq. (1) and (2)), initiating the communities near the given fixed point. We find that all species survive together until *E*(*G, C*) reaches 1, after which the mean survival fraction decreases approximately linearly with the encroachment (the slope is discussed in SI [42]). These results provide further support for our stability criterion, and indicate that encroachment is indicative of community behavior beyond the instability transition. If we imagine consumption ranges are constant and the reason for large encroachment is too small niche ranges due to large *N*_*S*_, the part of encroachment exceeding 1 in turn indicates how many species should be removed to restore stability and thus is related to the survival fraction. An analysis of the slope suggests that on average two species go extinct for each unstable eigenmode of the Jacobian (Fig. S1 in SI [42]). We therefore find that encroachment robustly predicts species loss after the transition to instability.

We then explored all the stable states and dynamics available beyond just perturbing the fixed point. To do so, we initialized communities by randomly sampling species abundances and resource concentrations and then quantified the diversity and dynamics of the final attractor. The fraction of survival shows the same trend (Fig. 4A) as the result obtained from perturbing the fixed point (Fig. 3E), with survival fraction not decreasing obviously until *E*(*G, C*) reaches 1. We found that fluctuating communities have higher diversity than stable communities with similar conditions, i.e., the same encroachment value *E*(*G, C*). Once encroachment *E*(*G, C*) surpasses 1, we observe the emergence of alternative stable states, globally stable states with species extinction, and fluctuating attractors (Fig. 4B; fluctuations include both limit cycles and chaos shown in Fig. S4 [42]). With increasing encroachment *E*(*G, C*), the fraction of fully coexisting globally stable states gradually decreases, the frequency of reaching alternative stable states increases, and observations of fluctuations peak at intermediate encroachment values (Fig 4B). Other resource supply methods may have different detailed behaviors but share the same basic trend (see Fig. S4 [42]). Encroachment *E*(*G, C*) as a measure of how strong species are encroaching others’ niches not only captures instability but also can organize seemingly scattered dynamical behaviors.

**Fig. 4.**
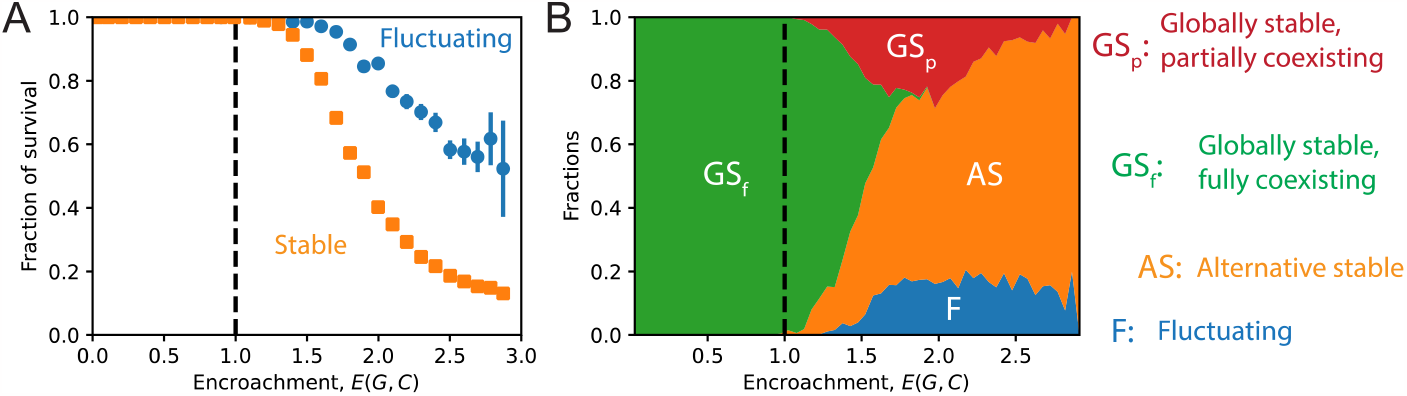
Beyond fixed points, various dynamical behaviors depend on encroachment. (A) Fluctuating communities have higher diversity under similar conditions. (B) Fractions of four different dynamical behaviors as a function of encroachment: GS_f_, i.e., the fully coexisting fixed point is globally stable, AS, i.e., alternative stable states, F, i.e., fluctuation, and GS_p_, i.e., a boundary fixed point is globally stable. Simulation results are obtained from communities with *N*_*S*=_16, *N*_*R*=_32, and boundaries between different regions in (B) are obtained from mean values [42]. Error bars in (A) represent SEM.

The community level behaviors also depend on how many resources and species are interacting. By varying the number of species *N*_*S*_ from 4 to 48 we calculated phase diagrams describing how the species-resource ratio *N*_*S*_ /*N*_*R*_ and encroachment *E*(*G, C*) alters the fractions of surviving species, fluctuating communities, and multistable communities (Fig. 5). By varying the number of resources we found that the species-resource ratio *N*_*S*_ /*N*_*R*_ is a characteristic variable that affect statistical behaviors of a complex community (various phase diagrams with different conditions presented in Fig. S5-S7 [42]). The existence of clear boundaries in all three phase diagrams indicates that encroachment is capturing essential information of the dynamics of communities. We found that for small *N*_*S*_ /*N*_*R*_ the onset of species extinction and community fluctuation occurs for encroachment *E*(*G, C*) > 1, consistent with previous claims since *E*(*G, C*) > 1 is a necessary but not sufficient condition of instability. To explain the *N*_*S*_ /*N*_*R*_ dependence of instability boundary, we first need to understand that encroachment *E*(*G, C*) > 1 merely enables a (small) probability for a pair of species to be unstable. When *N*_*S*_ /*N*_*R*_ increases, there are more pairs, increasing the probability of instability (SI [42]), and hence explaining why *E*(*G, C*) > 1 was a sufficient condition for predicting instability in previously (Fig 2 and 3). Based on this simple probability argument, we can derive a semi-empirical scaling law well describing the boundary of instability (line in Fig. 5). Alternative stable states arise earlier than losing local stability of the given fixed point, especially when *N*_*S*_ /*N*_*R*_ > 1, which can be partly explained from the perspective that *N*_*S*_ /*N*_*R*_ > 1, can lead to, in principle, infinite fixed-point solutions for the species abundances. In summary, by varying encroachment and the number of species we can we obtain the full phase diagrams related to stability and dynamics, highlighting that *E*(*G, C*) and *N*_*S*_ /*N*_*R*_ are the key determinants of community behaviors.

**Fig. 5.**
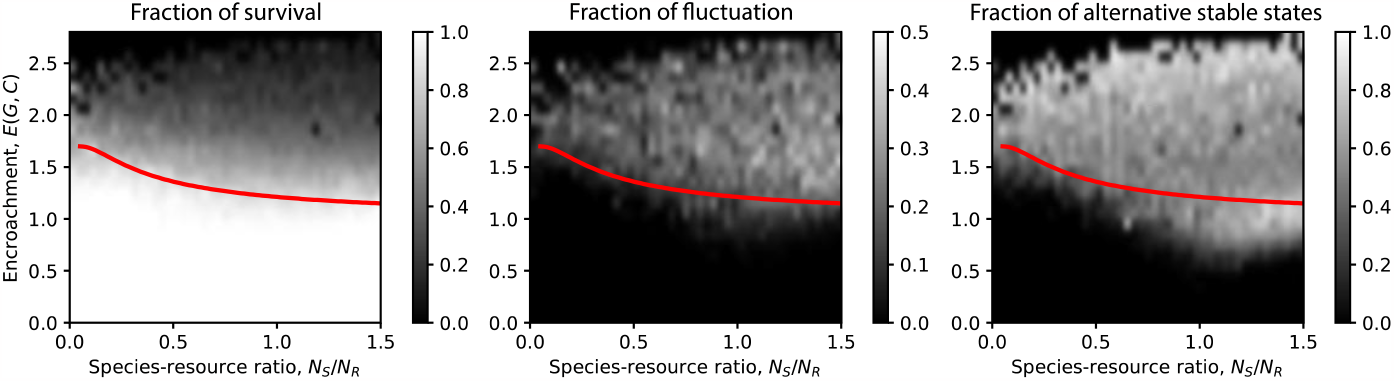
Encroachment and species-resource ratio, *N*_*S*_ /*N*_*R*_, shape phases of community diversity and dynamics. Boundary of being locally stable does not faithfully obey *E*(*G, C*)=1 for small *N*_*S*_ /*N*_*R*_ which can be predicted by a scaling based on probability argument [42]. Fraction of survival does not decrease clearly before losing local stability where there is almost no fluctuation, while alternative stable states can arise. Simulations are done for communities with *N*_*R*=_32 and *N*_*S*_ ranging from 4 to 48, and are initialized randomly. The color maps typically depict the mean value over more than 1000 simulations.

## Discussion

The mechanisms driving the intricate dynamics in natural communities are key to understanding and engineering ecological systems yet are not well understood. In this work, we studied the possible dynamics and diversity loss in communities driven by resource competition. We found that a discrepancy between resource consumption and species growth can lead to the full range of dynamics even in the simplest resource-consumer framework, consistent with recent reports [47]. Starting with local stability, we developed a local information free measure, the encroachment *E*(*G, C*), which captures many aspects of community dynamics. Our definition of consumption ranges, niche ranges, and their ratio, the encroachment have clear mechanistic meaning (Fig 2). We find that local stability is typically independent of species abundances and resource concentrations at the fixed point (Fig. 3). Encroachment—as a measure of how far away species are consuming from their own niches—not only predicts stability, but also yields survival fraction after the instability transition and provides insights regarding the complicated dynamical behaviors in this phase (Fig. 4). As discussed, consuming out of niche ranges, i.e., *E*(*G, C*) > 1, is a necessary but not sufficient condition of instability. However, when the number of species is sufficiently large, one’s neighboring niches are all occupied and configurations leading to instability will almost certainly be present (Fig. 5). The characterization of encroachment can help to understand dynamical behaviors of natural communities and potentially enable us to alter systems towards desired states via controlling encroachment properties.

The present analysis unifies and generalizes mechanistic understanding from different methods. First of all, Tilman’s graphic method is generalized to complex systems, showing the power of simple geometric reasoning. Second, we clarify how niche overlap limits coexistence. The concept of niche overlap [44] quantifies the similarity between niches which has the same role of niche range here. Increasing niche overlap or decreasing niche range can indeed drive the system towards instability, but we demonstrate that predicting loss of stability requires a comparison between niche range and consumption range. Third, we present mechanistic insights for D-stability, a method that was used to demonstrate global stability in the original MacArthur model [36]. We can show that *E*(*G, C*)=*0* is equivalent to D-stability (see Sec. S-I G in SI [42]). Looking for the existence of certain positive diagonal matrix *D* making the Lyapunov function converge can be abstract. Here, we suggest that this action is equivalent to finding a virtual fixed point where the system is most likely to be stable (Fig. 3C). Fourth, our work helps to bridge random matrix approach and resource competition. Since our reduced interaction matrix from resource competition is not i.i.d., a naïve application of May’s result [16] does not perform well (Fig. 2E). Instead, we show that, by using the correct theory for *GC*^*T*^, the simplified Jacobian, we can obtain a new stability criterion (Eq. (6)). This criterion recapitulates how the relationship between the consumption range and niche range determines stability of an ecosystem. Finally, our theory implies that besides the conventional competitive exclusion principle [34,35] based on the limit of feasibility, there is a new one based on stability. With a fixed consumption range, there cannot be too many species in a stable community otherwise the niche range will be too small leading to undesired encroachment. With the development of encroachment, we are able to obtain a picture unifying previous results.

Finally, new questions arise from the present work, which are not only related to resource competing communities and ecology but also to other complex systems. Our observation that local stability is independent of the specific location of fixed points needs further investigation. Existing random matrix analysis [45,48] has justified such observations in other complex systems but may not suffice here due to the complex emergent correlations in resource competition. In this work, we assumed the existence of a feasible fixed point to focus on local stability, which should be relaxed in future studies. Feasibility can be probed via cavity calculation which can further reveal the transition from single attractor phase to multiple attractors phase representing the loss of global stability [18,47]. As a comparison, cavity calculation [18,47] does not assume fixed points while encroachment analysis needs fewer assumptions about correlations within *G* or *C* (we only need *G*_*i*_ to be independent of *C*_*j*_ for *i* ≠ *j*, which is crucial in deriving the scaling in Fig. 5), but results from both methods (e.g., feasibility, global stability, and local stability) complement each other to provide a comprehensive understanding of various dynamic phases in general situations. Further research is needed to integrate the findings on feasibility with the new insights on stability gained here through the study of encroachment.

## Methods

We use Julia to conduct numerical tests and run simulations and use Python for data analysis and drawing figures. Simulations are run on MIT Supercloud [49]. All codes are available at https://github.com/liuyz0/RCM-stability. Usually, there are several modules in each simulation process: 1. Determining system size: We need to first specify the number of species and resources at the given fixed point, i.e., *N*_*S*_ and *N*_*R*_ ; 2. Sampling: Given *N*_*S*_ and *N*_*R*_, we then sample different consumption rates *C*, as well as growth rates *G*. The fixed point, embodied by equilibrium species abundances *S*^*\**^ and equilibrium resource concentrations *R*^*\**^ will also be sampled; 3. Simulating: With the sampled parameters, ordinary differential equations, Eqs. (1) and (2), are simulated with various initial conditions (typically one experiment involves 0.6 million simulation cases); 4. Analyzing data: A part of the data, for instance, the Jacobian, the characteristic matrix *GC*^*T*^, and the encroachment, can be directly calculated and analyzed at once after sampling. Other results, like fraction of survival, are obtained during dynamics simulations.

In the paper, there are two sampling ways, one is with strict metabolic trade-off ensuring uniform equilibrium resource concentrations (special sampling, used in Fig. 2), and the other samples everything including the fixed point randomly with uniform distributions (general sampling, used after Fig.2). The metabolic trade-off requires growth vectors *G*_*i*_ to be on the simplex. To obtain most diverse growth vectors, we sample *G*_*i*_ uniformly from the simplex. More discussions about sampling in each figure and related mathematical realizations can be found in Sec. S-II A of SI [42].

In Julia, we use the DifferentialEquations package and its solvers. Most times, the dynamics are non-stiff, and we tried solvers like Tsit5 (Tsitouras 5/4 Runge-Kutta method), Vern7 (Verner’s “Most Efficient” 7/6 Runge-Kutta method), and VCABM3 (the 3rd order Adams-Moulton method), which have no obvious differences in results. However, for logistic resource supply, the dynamics can be stiff and may require stiff solvers like Rodas4 (4th order A-stable stiffly stable Rosenbrock method with a stiff-aware 3rd order interpolant) and TRBDF2 (a second order A-B-L-S-stable one-step ESDIRK method). A callback function is needed to force the solutions to be non-negative, preventing non-physical results and maintaining numerical stability. Detailed discussions about solvers, solver parameters (e.g., checking time points for verifying convergence), and observable outputs are in SI (see Sec. S-II B [42]).

For data analysis, most quantities can be directly and simply calculated based on definitions. The calculation of *E*(*G, C*) is a little different since optimization is involved. We used PyTorch in python or Flux in Julia to write self-defined layer to output *CD* with *C* as the input and *D* the parameters to optimize. The loss function is given by *E*_*s*_(*G, C*). We tried different solvers like Adam and stochastic gradient descent (SGD) which yield the same results. The learning rate is set as 0.1 and the number of iterations is 1000. These settings have been tested to guarantee convergence for all optimization processes.

Theoretical reasoning leading to our main messages is presented in Results. Other analysis methods, which is related but not necessary in the path to encroachment, like Tilman’s original graphic analysis, D-stability (Lyapunov function), local stability analysis (Jacobian), and random matrix results, can be found in previous literatures and for completeness are reviewed and collected in SI.

## Acknowledgments

We thank all the Gore Laboratory members for their valuable suggestions. We are grateful for Akshit Goyal’s help in understanding resource-consumer models. We appreciate Prof. Pankaj Mehta and his group members at Boston University for inspiring and fruitful discussions. We thank Matthieu Barbier and Prof. Guy Bunin for their constructive suggestions which improves the manuscript a lot. J.G. thanks the Schmidt Foundation and Sloan Foundation for funding.

## References

[1] J. J. Faith et al., The Long-Term Stability of the Human Gut Microbiota, Science 341, 1237439 (2013).

[2] D. Tilman, P. B. Reich, and J. M. H. Knops, Biodiversity and Ecosystem Stability in a Decade-Long Grassland Experiment, Nature 441, 7093 (2006).

[3] M. Dodd, J. Silvertown, K. McConway, J. Potts, and M. Crawley, Community Stability: A 60-Year Record of Trends and Outbreaks in the Occurrence of Species in the Park Grass Experiment, Journal of Ecology 83, 277 (1995).

[4] A. Schröder, L. Persson, and A. M. De Roos, Direct Experimental Evidence for Alternative Stable States: A Review, Oikos 110, 3 (2005).

[5] M. Van de Guchte, S. D. Burz, J. Cadiou, J. Wu, S. Mondot, H. M. Blottière, and J. Doré, Alternative Stable States in the Intestinal Ecosystem: Proof of Concept in a Rat Model and a Perspective of Therapeutic Implications, Microbiome 8, 153 (2020).

[6] C. Averill, C. Fortunel, D. S. Maynard, J. van den Hoogen, M. C. Dietze, J. M. Bhatnagar, and T. W. Crowther, Alternative Stable States of the Forest Mycobiome Are Maintained through Positive Feedbacks, Nat Ecol Evol 6, 4 (2022).

[7] J. van de Koppel, P. M. J. Herman, P. Thoolen, and C. H. R. Heip, Do Alternate Stable States Occur in Natural Ecosystems? Evidence from a Tidal Flat, Ecology 82, 3449 (2001).

[8] D. R. Amor, C. Ratzke, and J. Gore, Transient Invaders Can Induce Shifts between Alternative Stable States of Microbial Communities, Science Advances 6, eaay8676 (2020).

[9] C. I. Abreu, V. L. A. Woltz, J. Friedman, and J. Gore, Microbial Communities Display Alternative Stable States in a Fluctuating Environment, PLOS Computational Biology 16, e1007934 (2020).

[10] B. Blasius, L. Rudolf, G. Weithoff, U. Gaedke, and G. F. Fussmann, Long-Term Cyclic Persistence in an Experimental Predator–Prey System, Nature 577, 7789 (2020).

[11] F. K. Balagaddé, H. Song, J. Ozaki, C. H. Collins, M. Barnet, F. H. Arnold, S. R. Quake, and L. You, A Synthetic Escherichia Coli Predator–Prey Ecosystem, Molecular Systems Biology 4, 187 (2008).

[12] G. F. Fussmann, S. P. Ellner, K. W. Shertzer, and N. G. Hairston Jr., Crossing the Hopf Bifurcation in a Live Predator-Prey System, Science 290, 1358 (2000).

[13] E. Benincà, B. Ballantine, S. P. Ellner, and J. Huisman, Species Fluctuations Sustained by a Cyclic Succession at the Edge of Chaos, Proceedings of the National Academy of Sciences 112, 6389 (2015).

[14] E. Benincà, J. Huisman, R. Heerkloss, K. D. Jöhnk, P. Branco, E. H. Van Nes, M. Scheffer, and S. P. Ellner, Chaos in a Long-Term Experiment with a Plankton Community, Nature 451, 7180 (2008).

[15] C. Ratzke, J. Barrere, and J. Gore, Strength of Species Interactions Determines Biodiversity and Stability in Microbial Communities, Nat Ecol Evol 4, 3 (2020).

[16] R. M. May, Will a Large Complex System Be Stable?, Nature 238, 5364 (1972).

[17] S. Allesina and S. Tang, Stability Criteria for Complex Ecosystems, Nature 483, 7388 (2012).

[18] G. Bunin, Ecological Communities with Lotka-Volterra Dynamics, Phys. Rev. E 95, 042414 (2017).

[19] D. A. Kessler and N. M. Shnerb, Generalized Model of Island Biodiversity, Phys. Rev. E 91, 042705 (2015).

[20] J. Hu, D. R. Amor, M. Barbier, G. Bunin, and J. Gore, Emergent Phases of Ecological Diversity and Dynamics Mapped in Microcosms, Science 378, 85 (2022).

[21] A. Altieri, F. Roy, C. Cammarota, and G. Biroli, Properties of Equilibria and Glassy Phases of the Random Lotka-Volterra Model with Demographic Noise, Phys. Rev. Lett. 126, 258301 (2021).

[22] V. Ros, F. Roy, G. Biroli, G. Bunin, and A. M. Turner, Generalized Lotka-Volterra Equations with Random, Nonreciprocal Interactions: The Typical Number of Equilibria, Phys. Rev. Lett. 130, 257401 (2023).

[23] F. Roy, M. Barbier, G. Biroli, and G. Bunin, Complex Interactions Can Create Persistent Fluctuations in High-Diversity Ecosystems, PLOS Computational Biology 16, e1007827 (2020).

[24] T. A. de Pirey and G. Bunin, Many-Species Ecological Fluctuations as a Jump Process from the Brink of Extinction, arXiv:2306.13634.

[25] M. T. Pearce, A. Agarwala, and D. S. Fisher, Stabilization of Extensive Fine-Scale Diversity by Ecologically Driven Spatiotemporal Chaos, Proceedings of the National Academy of Sciences 117, 14572 (2020).

[26] T. Arnoulx de Pirey and G. Bunin, Aging by Near-Extinctions in Many-Variable Interacting Populations, Phys. Rev. Lett. 130, 098401 (2023).

[27] M. Dal Bello, H. Lee, A. Goyal, and J. Gore, Resource–Diversity Relationships in Bacterial Communities Reflect the Network Structure of Microbial Metabolism, Nat Ecol Evol 5, 10 (2021).

[28] P.-Y. Ho, B. H. Good, and K. C. Huang, Competition for Fluctuating Resources Reproduces Statistics of Species Abundance over Time across Wide-Ranging Microbiotas, eLife 11, e75168 (2022).

[29] P.-Y. Ho, T. H. Nguyen, J. M. Sanchez, B. C. DeFelice, and K. C. Huang, Resource Competition Predicts Assembly of in Vitro Gut Bacterial Communities.

[30] A. Posfai, T. Taillefumier, and N. S. Wingreen, Metabolic Trade-Offs Promote Diversity in a Model Ecosystem, Phys. Rev. Lett. 118, 028103 (2017).

[31] M. Tikhonov and R. Monasson, Collective Phase in Resource Competition in a Highly Diverse Ecosystem, Phys. Rev. Lett. 118, 048103 (2017).

[32] W. Cui, R. Marsland, and P. Mehta, Effect of Resource Dynamics on Species Packing in Diverse Ecosystems, Phys. Rev. Lett. 125, 048101 (2020).

[33] E. Blumenthal and P. Mehta, Geometry of Ecological Coexistence and Niche Differentiation.

[34] G. Hardin, The Competitive Exclusion Principle, Science 131, 1292 (1960).

[35] S. A. Levin, Community Equilibria and Stability, and an Extension of the Competitive Exclusion Principle, The American Naturalist 104, 413 (1970).

[36] R. M. Arthur, Species Packing, and What Competition Minimizes*†, Proceedings of the National Academy of Sciences 64, 1369 (1969).

[37] S. Butler and J. P. O’Dwyer, Stability Criteria for Complex Microbial Communities, Nat Commun 9, 1 (2018).

[38] I. Dalmedigos and G. Bunin, Dynamical Persistence in High-Diversity Resource-Consumer Communities, PLOS Computational Biology 16, e1008189 (2020).

[39] J. Huisman and F. J. Weissing, Biological Conditions for Oscillations and Chaos Generated by Multispecies Competition, Ecology 82, 2682 (2001).

[40] B. Bloxham, H. Lee, and J. Gore, Diauxic Lags Explain Unexpected Coexistence in Multi-Resource Environments, Molecular Systems Biology 18, e10630 (2022).

[41] D. Tilman, Resource Competition and Community Structure (Princeton University Press, 1982).

[42] Materials and Methods Are Available in Supplementary Information., (n.d.).

[43] A. D. Letten, P.-J. Ke, and T. Fukami, Linking Modern Coexistence Theory and Contemporary Niche Theory, Ecological Monographs 87, 161 (2017).

[44] R. M. May and R. H. M. Arthur, Niche Overlap as a Function of Environmental Variability, Proceedings of the National Academy of Sciences 69, 1109 (1972).

[45] L. Stone, The Feasibility and Stability of Large Complex Biological Networks: A Random Matrix Approach, Sci Rep 8, 1 (2018).

[46] Vinayak and L. Benet, Spectral Domain of Large Nonsymmetric Correlated Wishart Matrices, Phys. Rev. E 90, 042109 (2014).

[47] E. Blumenthal, J. W. Rocks, and P. Mehta, Phase Transition to Chaos in Complex Ecosystems with Non-Reciprocal Species-Resource Interactions, arXiv:2308.15757.

[48] Y. Ahmadian, F. Fumarola, and K. D. Miller, Properties of Networks with Partially Structured and Partially Random Connectivity, Phys. Rev. E 91, 012820 (2015).

[49] A. Reuther et al., Interactive Supercomputing on 40,000 Cores for Machine Learning and Data Analysis, in 2018 IEEE High Performance Extreme Computing Conference (HPEC) (2018), pp. 1–6.

