## Supplemental Information for "Complex ecosystems lose stability when resource consumption is out of niche"

### Supplementary information for "Complex ecosystems lose stability when resource consumption is out of niche"

#### CONTENTS

S-I. Theory 1
A. Revisiting the models 1
B. Tilman analysis 2
C. Local Jacobian 3
D. Encroachment and stability criteria 4
E. Interaction matrix measure 5
F. Characteristic matrix 6
G. Global stability 6

S-II. Numerical methods 7
A. Sampling methods 8
B. Simulation methods 8
C. Data analysis methods 9

S-III. Figure details and supplementary results 9
A. Figure 1 10
B. Figure 2 10
C. Figure 3 11
D. Figure 4 12
E. Figure 5 12

References 13

#### S-I. THEORY

##### A. Revisiting the models

As a reminder, the models analyzed and the related definitions are reviewed. Dynamics between species $S_i$ and resources $R_\alpha$ is governed by

$$\frac{\mathrm{d}S_i}{\mathrm{d}t} = S_i\Big(\sum_\alpha G_{i\alpha}R_\alpha - \delta_i\Big), \tag{S1a}$$

$$\frac{\mathrm{d}R_\alpha}{\mathrm{d}t} = h_\alpha(R_\alpha) - R_\alpha \sum_i C_{i\alpha}S_i, \tag{S1b}$$

where $G_{i\alpha}$ is is the per capital growth rate of species $i$ with unit resource $\alpha$, $\delta_i$ is the mortality of species $i$, $h_\alpha(R_\alpha)$ encodes how resources are supplied, and $C_{i\alpha}$ denotes the per capital consumption rate of species $i$ for resource $\alpha$. Species consume different resources to grow which reduces resource concentrations, while species are also decreasing subject to mortality and resources are supplied into the system.

---

*
†

Different resource supply functions $h_\alpha(R_\alpha)$ correspond to different types of resources. In the earliest works on resource-consumer models [S1], biotic resources are considered in the context of zoology, resulting a logistic form, $h_\alpha(R_\alpha) = g_\alpha R_\alpha(K_\alpha - R_\alpha)$. In chemostats, resources flow in from certain source, giving a linear function $h_\alpha(R_\alpha) = l_\alpha(\kappa_\alpha - R_\alpha)$. In the main text, we present the simplest constant form $h_\alpha(R_\alpha) = \gamma_\alpha$, which can be taken as an approximation of other resource supply methods.

##### B. Tilman analysis

For completeness, the original Tilman's graphical analysis is reviewed [S2]. In the main text, we only present key information to determine stability without detailed explanations. Here, we will show the full picture, not only about stability but also feasibility, providing concrete understating for simple communities with two species and two resources. Tilman made use of the graphs in resource (Fig. S2), where each axis represents one resource concentration. We use the red axis for resource 1, and blue one for resource 2. By solving the equation

$$\frac{\mathrm{d}S_1}{\mathrm{d}t} = \sum_{\alpha=1}^{2} G_{1\alpha} R_\alpha - \delta_1 = 0, \tag{S2}$$

we can get one line in resource concentration space (the red solid line in Fig. S2), on which species 1 has no net growth. Such a line is called a zero net growth isocline (ZNGI). Similarly, we can have another ZNGI of species 2 (the blue solid line inside patches of Fig. S2) by solving

$$\frac{\mathrm{d}S_2}{\mathrm{d}t} = \sum_{\alpha=1}^{2} G_{2\alpha} R_\alpha - \delta_2 = 0. \tag{S3}$$

Existence of a feasible coexisting fixed point depends on the ZNGIs, resource supply, and consumption. First, if we want the two resources to coexist at certain steady concentrations where species have zero net growth, the two ZNGIs will need to have a intersection in the region $R_1,\ R_2 > 0$ (e.g., Fig. S2). The intersection of ZNGIs, $R^* = (R_1^*, R_2^*)$, encode resource concentrations making species abundances steady. Second, in making the species to coexist at steady and positive abundances, the following is necessary:

$$\frac{\mathrm{d}R_\alpha}{\mathrm{d}t} = h_\alpha(R_\alpha^*) - R_\alpha^* \sum_{i=1}^{2} C_{i\alpha} S_i^* = 0, \text{ for } \alpha = 1, 2, \tag{S4}$$

from which we can solve the equilibrium species abundances $S^*$. In the matrix form, we can define $V = CD(R^*)$ where $D(v)$ denotes a diagonal matrix with vector $v$ on the diagonal. The impact vector of species $i$, $V_i$, is the $i$th row of the matrix $V$, encoding what "direction" of resources is consumed by species $i$ at the fixed point. Rewriting Eq. (S4) with vectors, we have

$$h(R^*) - S_1^* V_1 - S_2^* V_2 = 0, \tag{S5}$$

which suggests that $h(R^*)$ is a linear combination of $V_1$ and $V_2$. In the graph, resource supply $h(R^*)$ can be denoted by a vector starting from the intersection of ZNGIs (black arrow in the left patch of Fig. S2), and so do the impact vectors (the direction of $V_1$ ($V_2$) is shown by the red (blue) dashed line in Fig. S2). The requirement that species abundances at the fixed point, $S^*$, is positive, is equivalent to requiring $h(R^*)$ to be between the impact vectors (for example, being in the yellow region of the left patch, or in the gray region of the right patch of Fig. S2). For a coexisting feasible fixed point to exist, we need the ZNGIs to intersect at a point with all positive concentrations giving $R^*$, and need the supply vector $h(R^*)$ to be between the impact vectors giving feasible $S^*$. If either of the two conditions is not satisfied, only one species can survive. If in the all positive quadrant, ZNGIs do not intersect, species correspond to the ZNGI closer to the zero point (i.e., species with higher growth rates) will survive. If we have feasible $R^*$ but $h(R^*)$ is not between the impact vectors, the species whose impact vector is loser to the supply vector will survive (for instance, in the left patch of Fig. S2, if the black arrow is in the blue (red) region, only species 2 (1) can survive). In a word, Tilman's graph can clearly show the conditions of having a coexisting fixed point, as well as which species can survive once feasibility is lost.

Given the existence of the coexisting fixed point, Tilman's graph can determine whether the fixed point is locally stable or not. To consider stability, the growth vector $G_i$, being the $i$th row of matrix $G$ is needed besides impact vectors. From Eqs. (S2)-(S3) which define ZNGIs, we can know that the vector $G_i$ is orthogonal to the ZNGI of species $i$. The case $V_i \parallel G_i$ is heuristically considered as most stable (left patch of Fig. S2, where the squares show directions of the growth vectors starting from the intersection), since the species tries to drag resources back to equilibrium by consumption in the rapidest direction (normal direction of the ZNGI). In fact, as long as one's impact vector is more aligned with the growth vector of itself than the other one's impact vector, the fixed point will be stable. The idea can be simply explained with feedback loops. Assume that species 1 grows better on resource 1 than species 2. If species 1 consume more resource 1 ($V_i$ aligned more with $G_i$), increasing resource 1 a little above equilibrium, species 1 will grow better due to the direction of growth vector and resource 1 will be rapidly consumed and decrease due to consumption from species 1. Let impact vectors be closer to the other sides' growth vectors, the critical situation happens when the two impact vectors have the same direction, leading to marginal stability (the middle patch of Fig. S2). After that, if species 1 consume resource 2 better ($V_i$ aligned more with $G_j$), still increasing resource 1 a little above equilibrium, species 1 will grow faster, resource 2 will be decreased more (by consumption), species 2 then will decrease (due to growth preference), and resource 1 can grow more without its major consumer, resulting a positive feedback loop and thus instability (right patch of Fig. S2 shows a special example of instability). In graphic representations, the instability transition is vividly shown as flipping of impact vectors, since the two lines representing vectors will collapse and change the order of which one is above the other.

Geometric understanding from Tilman's graph has algebraic representations, revealing local stability is irrelevant to fixed-point information. The intuition of flipping the two dimensional impact vectors can be rigorously described via cross product. Consider the products,

$$\begin{pmatrix} G_{11} \\ G_{12} \\ 0 \end{pmatrix} \times \begin{pmatrix} G_{21} \\ G_{22} \\ 0 \end{pmatrix} \propto \begin{pmatrix} 0 \\ 0 \\ \sin\theta_G \end{pmatrix}, \tag{S6}$$

and

$$\begin{pmatrix} V_{11} \\ V_{12} \\ 0 \end{pmatrix} \times \begin{pmatrix} V_{21} \\ V_{22} \\ 0 \end{pmatrix} \propto \begin{pmatrix} 0 \\ 0 \\ \sin\theta_V \end{pmatrix}, \tag{S7}$$

where $\theta_G$ is the angle from vector $G_1$ to $G_2$, and $\theta_V$ the angle from vector $V_1$ to $V_2$. As long as the two angles have the same sign, the stability holds as $V_i$ is more aligned with $G_i$. On the contrary, if the two angles have different signs, flipping of impact vectors occur, resulting instability. Recall the relation between cross product and determinant, e.g.,

$$\begin{vmatrix} V_{11} & V_{12} \\ V_{21} & V_{22} \end{vmatrix} \propto \sin\theta_V, \tag{S8}$$

we can write down the stability criterion as

$$\begin{vmatrix} V_{11} & V_{12} \\ V_{21} & V_{22} \end{vmatrix} \begin{vmatrix} G_{11} & G_{12} \\ G_{21} & G_{22} \end{vmatrix} > 0, \tag{S9}$$

or simply,

$$\det(GV) > 0, \text{ or } \det(GV^\top) > 0 \tag{S10}$$

Note that $V = CD(R^*)$, we have $\det(GV) = \det(GC)\det(D(R^*))$. And since $R^*$ are all positive, it does not change the sign, such that, the above stability criterion is equivalent to

$$\det(GC) > 0, \text{ or } \det(GC^\top) > 0. \tag{S11}$$

If the number of species is not the same as that of resources, $GC$ does not exist. Therefore, the form $GC^\top$ is the generalizable one. The algebraic reasoning helps us to see the fact that local information like $R^*$ does not affect stability and gives the simple form $GC^\top$, which inspires the refined encroachment $E(G, C)$ (Fig. 3).

##### C. Local Jacobian

Local stability is canonically detected via local Jacobian analysis, which is mathematically introduced and studied here. Local Jacobian is defined at the fixed point of interest (specified by equilibrium abundances $S^*$ and concentrations $R^*$) via linearization of the dynamics. Let $s_i$ and $r_\alpha$ be perturbations of species and resources, i.e., $s_i = S_i - S_i^*$

and $r_\alpha = R_\alpha - R_\alpha^*$, by assuming the perturbations are small, we have the dynamics reduced from Eq. (S1) only keeping first order terms:

$$\frac{\mathrm{d}}{\mathrm{d}t}\begin{pmatrix} s_1 \\ \vdots \\ s_{N_S} \\ \hline r_1 \\ \vdots \\ r_{N_R} \end{pmatrix} = \left(\begin{array}{c|c} O & D(S^*)G \\ \hline -D(R^*)C^\top & \frac{\partial h}{\partial R}|_{R^*} - D(C^\top S^*) \end{array}\right)\begin{pmatrix} s_1 \\ \vdots \\ s_{N_S} \\ \hline r_1 \\ \vdots \\ r_{N_R} \end{pmatrix}, \tag{S12}$$

where $N_S$ and $N_R$ are the number of surviving species and resources at the fixed point, respectively (we do not consider invasion of species from external pools). The local Jacobian is defined as

$$J^* = \left(\begin{array}{c|c} O & D(S^*)G \\ \hline -D(R^*)C^\top & \frac{\partial h}{\partial R}|_{R^*} - D(C^\top S^*) \end{array}\right). \tag{S13}$$

The $N_R \times N_R$ matrix $\frac{\partial h}{\partial R}$ has elements $\frac{\partial h_\alpha}{\partial R_\beta}$. For constant supply, $h_\alpha = \gamma_\alpha$, $\frac{\partial h}{\partial R} = 0$; for linear supply, $h_\alpha = l_\alpha(\kappa_\alpha - R_\alpha)$, $\frac{\partial h}{\partial R} = -D(l)$; and for logistic supply, $h_\alpha = g_\alpha R_\alpha(K_\alpha - R_\alpha)$, diagonal elements are $\frac{\partial h_\alpha}{\partial R_\alpha} = g_\alpha(K_\alpha - 2R_\alpha)$ and off-diagonal ones are zero. If all the eigenvalues of the Jacobian have negative real parts, the perturbations will vanish exponentially, suggesting stability. On the contrary, if there exists any eigenvalue with positive real part, the perturbation can grow exponentially, leading to instability. In the paper (e.g., Fig. 2 and Fig. 3), we numerically solve the eigenvalues of the Jacobian to determine stability of the given fixed point of interest.

Note that at the fixed point

$$\frac{\mathrm{d}R}{\mathrm{d}t} = h(R^*) - D(R^*)D(C^\top S^*) = 0, \tag{S14}$$

it is easy to verify that for all three resource supply methods of interest, the diagonal matrix in the lower right part of $J^*$

$$\left.\frac{\partial h}{\partial R}\right|_{R^*} - D(C^\top S^*) = \begin{cases} -D(C^\top S^*) & \text{Constant supply,} \\ -D(l) - D(C^\top S^*) & \text{Linear supply,} \\ -D(g)D(R^*) & \text{Logistic supply,} \end{cases} \tag{S15}$$

can only have negative elements.

The eigenvalues of the Jacobian can be solved via $\det(zI - J^*) = 0$ where $z$ is the variable and $I$ the identity matrix, which gives

$$\det\left(zI + D(S^*)GD(R^*)\left(zI - \left.\frac{\partial h}{\partial R}\right|_{R^*} + D(C^\top S^*)\right)^{-1} C^\top\right)\det\Big(zI - \left.\frac{\partial h}{\partial R}\right|_{R^*} + D(C^\top S^*)\Big) = 0. \tag{S16}$$

##### D. Encroachment and stability criteria

We then discuss some subtle understanding of encroachment based on its relation with Jacobian. Recall the definition of encroachment is the mean ratio of consumption ranges and niche ranges. When encroachment, the mean value, is smaller than 1, there can be some outliers or extreme value exceeding 1, possibly leading to instability. In the limit that $N_S$ goes to infinity, the probability of having such outliers given the mean value, encroachment, is smaller than 1, will tend to be 1. This means that encroachment will fail to identify the absolute stability when system size is extremely large due to the fact it is just an average value. However, in the large $N_S$ limit, if we only have a few outliers, although the system may be absolutely unstable and go to another state after extinction of certain outliers, the new state would be almost the same as the original one since the number of outliers is too small to make statistically significant changes to the whole system. The change due to instability will be significant, when the number of outliers is comparable to the system size. And the encroachment, as a mean value, certainly captures the behavior of a portion of individuals comparable to the total size, thus being a robust measure of statistical instability.

In terms of local Jacobian, we can define its fraction of unstable modes as

$$\frac{\#\text{ of eigenvalues with positive real parts}}{N_S}, \tag{S17}$$

which describes the degree of instability. Note that the denominator is $N_S$ since there are at most $N_S$ eigenvalues of the local Jacobian can have positive real parts. If the Jacobian has 1 unstable mode, the system is absolutely unstable. But when $N_S \to \infty$, its effect can be negligible. The system is statistically unstable when the fraction of unstable modes is non-negligible, i.e., the number unstable modes is comparable to the systems size. We show that the encroachment indeed strongly correlates with the fraction of unstable modes (Fig. S1).

Although the encroachment, as a mean value, is limited, it is possible to go beyond some limitations following our method with more details. We can write encroachment as $E = \text{mean}_i E_i$, where $E_i$ is the ratio of consumption range and niche range for species $i$. Assume $E_i$ are independent and identically distributed (i.i.d.), knowing the distribution of $E_i$ can calculate $P(E_i > 1)$ can show the possibility of absolute instability. Or, in a coarse-grained picture, we can make use of the mean value $E$ and the standard deviation $\text{std}(E_i)$ to estimate $P(E_i > 1)$. The statistical instability is related to whether $P(E_i > 1)$ is larger than certain fixed number. Knowing the standard deviation $\text{std}(E_i)$, we can also estimate how sharp the instability transition can be. The smaller the $\text{std}(E_i)$ is, the better the system is captured by encroachment alone, the smaller the window of encroachment will be where the fraction of unstable communities goes from 0 to 1.

##### E. Interaction matrix measure

Local mapping of resource-consumer models to the well-known generalized Lotka-Volterra (GLV) model is introduced here, followed by the definition of local interaction matrix. The GLV equation describe interactions between species in a phenomenological way:

$$\frac{\mathrm{d}S_i}{\mathrm{d}t} = \frac{b_i}{k_i} S_i (k_i - \sum_{i,j} a_{ij} S_j), \tag{S18}$$

where $S_i$ still denotes the abundance of the species $i$, $b_i$ are the intrinsic growth rates, $k_i$ are the carrying capacities, and $a_{ij}$ is the interaction matrix whose diagonal elements are 1. Suppose there is a fixed point $S^*$, near the fixed point, the perturbation $s = S - S^*$ is governed by the local linear dynamics

$$\frac{\mathrm{d}s_i}{\mathrm{d}t} = -\frac{b_i}{k_i} S_i^* \sum_{i,j} a_{ij} s_j. \tag{S19}$$

The local Jacobian of GLV is given by

$$(J^*_{\mathrm{GLV}})_{ij} = -\frac{b_i}{k_i} S_i^* a_{ij}. \tag{S20}$$

And since $\frac{b_i}{k_i} S_i^*$ are all positive, the stability is determined by the interaction $a_{ij}$, i.e., whether $J^*_{\mathrm{GLV}}$ has eigenvalues of positive eigenvalues is the same as whether the matrix $-a_{ij}$ does [S3, S4].

The key of local mapping is to reduce the resource consumer models to a form like Eq. (S18), find the corresponding phenomenological interaction matrix, and apply random matrix analysis to that interaction matrix. Assume that metabolic reactions occur much faster than cell growth, we can set resource net flux $\mathrm{d}R/\mathrm{d}t = 0$ and focus on the relatively slower species dynamics. Near the fixed point, we can learn from Eq. (S12) that

$$\frac{\mathrm{d}r}{\mathrm{d}t} = -D(R^*)C^\top s + [\frac{\partial h}{\partial R}|_{R^*} - D(C^\top S^*)]r = 0, \tag{S21}$$

and

$$\frac{\mathrm{d}s}{\mathrm{d}t} = D(S^*)Gr. \tag{S22}$$

Resource perturbation $r$ can be solved from the first equation being a function of species perturbation $s$. Substitute the solved $r$ to the second equation, we obtain a closed species interaction equation,

$$\frac{\mathrm{d}s}{\mathrm{d}t} = -D(S^*)G[-\frac{\partial h}{\partial R}|_{R^*} + D(C^\top S^*)]^{-1} D(R^*)C^\top s. \tag{S23}$$

Comparing to the local GLV dynamics, we find the interaction matrix abstracted from resource competition is

$$G[-\frac{\partial h}{\partial R}|_{R^*} + D(C^\top S^*)]^{-1} D(R^*)C^\top. \tag{S24}$$

Note that for GLV dynamics, $a_{ii} = 1$, we also resale inter-species interactions by intra-species ones for the resource explicit model, and define the rescaled interaction matrix as

$$A = D(\sqrt{d(A)^{-1}})AD(\sqrt{d(A)^{-1}}), \tag{S25}$$

where $d(\cdot)$ extracts the diagonal elements of a matrix.

Given a $N \times N$ random matrix with 1 in diagonal and i.i.d. random variables having (positive) mean $\mu$ and standard deviation $\sigma$ as off-diagonal elements, the criterion of having no eigenvalues with negative real parts is given by [S4]

$$\frac{\sqrt{N}\sigma}{1-\mu} < 1. \tag{S26}$$

Note the equivalence between the Jacobian and the negative interaction matrix in identifying stability, stability should be preserved if

$$\frac{\sqrt{N_S}\sigma_A}{1-\mu_A} < 1, \tag{S27}$$

given the premise we trust the random matrix theory to work for the local resource-consumer dynamics, where $\sigma_A$ and $\mu_A$ are the standard deviation and mean of off-diagonal elements of $A$, respectively. To avoid singularity produced by $1-\mu_A$, we define the interaction matrix measure (used in Fig. 2) as

$$M_{\mathrm{RMT}} = \sqrt{N_S}\sigma_A + \mu_A. \tag{S28}$$

Clearly, stability is by theory equivalent to $M_{\mathrm{RMT}} < 1$, which has been shown not to work since the calculated $M_{\mathrm{RMT}} > 1$ for stable communities.

The failure of $M_{\mathrm{RMT}}$ may be caused by the fact that elements of $A$ are not i.i.d. For example, when $C = G$, the matrix $A$ is symmetric, i.e., $A_{ij} = A_{ji}$. Considering the correlation between $A_{ij}$ and $A_{ji}$ may not make the predictions better. For the case $C \approx G$, i.e., $A$ is symmetric, $\sqrt{N_S}\sigma_A$ in $M_{\mathrm{RMT}}$ should be refined to $2\sqrt{N_S}\sigma_A$, which will increase the calculated $M_{\mathrm{RMT}}$ further away from 1. But by the theory, $C \approx G$ should lead to stability, or $M_{\mathrm{RMT}} < 1$. The complicated correlations among various elements (not only between symmetric ones, like $A_{ij}$ and $A_{ji}$) may have non-negligible effects and making the simple and widely used random matrix predictions fail.

#### F. Characteristic matrix

Knowing local stability is irrelevant to fixed-point information, we use two (non-rigorous) methods to eliminate local information in the Jacobian to obtain and discuss the characteristic matrix $GC^\top$. The first way is to make use of the interaction matrix Eq. (S24) obtained depending on the assumption that resource dynamics reaches equilibrium much faster than species dynamics. Since we know $[-\frac{\partial h}{\partial R}|_{R^*} + D(C^\top S^*)]^{-1}D(R^*)$ is a positive definite diagonal matrix. If we assume it does not affect the signs of eigenvalues, whether the local interaction matrix has unstable eigenmode is equivalent to whether $GC^\top$ does. The second method starts from the rigorous Eq. (S16) which the eigenvalues of Jacobian $J^*$ should satisfy. If we solve eigenvalues from

$$\det\left(zI + D(S^*)GD(R^*)\left(zI - \left.\frac{\partial h}{\partial R}\right|_{R^*} + D(C^\top S^*)\right)^{-1} C^\top\right) = 0, \tag{S29}$$

and still assume the diagonal positive matrices do not affect signs of eigenvalues, we can conclude that once $GC^\top$ has a positive (negative) real part eigenvalue, the Jacobian $J^*$ should also have a positive (negative) one.

#### G. Global stability

A sufficient condition for global stability can be given for logistic resource supply model (i.e., the MacArthur model [S1]). Rewriting the resource model Eq. (S1) with $h_\alpha(R_\alpha) = g_\alpha R_\alpha(K_\alpha - R_\alpha)$ in matrix form gives:

$$\frac{\mathrm{d}}{\mathrm{d}t}\begin{pmatrix} S \\ R \end{pmatrix} = \left(\begin{array}{c|c} D(S) & O \\ \hline O & D(R) \end{array}\right)\left[\left(\begin{array}{c|c} O & G \\ \hline -C^\top & -D(g) \end{array}\right)\begin{pmatrix} S \\ R \end{pmatrix} + \begin{pmatrix} -\delta \\ g \circ K \end{pmatrix}\right], \tag{S30}$$

where $(g \circ K)_\alpha = g_\alpha K_\alpha$ is a Hadamard product. By defining

$$x = \left(\frac{S}{R}\right),\ B = \left(\begin{array}{c|c} O & G \\ \hline -C^\top & -D(g) \end{array}\right), \tag{S31}$$

and introducing the fixed point $x^*$ (using $\frac{\mathrm{d}x}{\mathrm{d}t}|_{x^*} = 0$), we can obtain a simpler form of dynamics:

$$\frac{\mathrm{d}x}{\mathrm{d}t} = D(x)B(x - x^*). \tag{S32}$$

Construct a Lyapunov equation as

$$V(x) = \sum_i w_i(x_i - x_i^* \ln x_i), \tag{S33}$$

where $w_i > 0$ and $V(x)$ is bounded from below. Time derivative of $V(x)$ is given by

$$\begin{aligned} \frac{\mathrm{d}V(x)}{\mathrm{d}t} &= \sum_i w_i(\frac{\mathrm{d}x_i}{\mathrm{d}t} - \frac{x_i^*}{x_i}\frac{\mathrm{d}x_i}{\mathrm{d}t}) \\ &= \sum_{ij}(x_i - x_i^*)w_i B_{ij}(x_j - x_j^*) \\ &= (x - x^*)^\top \frac{1}{2}(D(w)B + B^\top D(w))(x - x^*). \end{aligned} \tag{S34}$$

If there exist a set of $w$, such that $D(w)B + B^\top D(w)$ is negative semi-definite, we can have $\frac{\mathrm{d}V(x)}{\mathrm{d}t} \le 0$, yielding global stability. Let the first $N_S$ elements of $w$ be a new vector $w_1$ and the left elements be $w_2$, we can expand $D(w)B + B^\top D(w)$ as

$$D(w)B + B^\top D(w) = \left(\begin{array}{c|c} O & D(w_1)G - CD(w_2) \\ \hline G^\top D(w_1) - D(w_2)C^\top & -2D(w_2)D(g) \end{array}\right). \tag{S35}$$

If $D(w_1)G - CD(w_2) = 0$, $D(w)B + B^\top D(w)$ is negative semi-definite. To conclude, if there exist positive vectors $w_1$ and $w_2$, such that $D(w_1)G - CD(w_2) = 0$, the fixed point $x^*$ is globally stable.

Next, we show that $E(G, C) = 0$ is equivalent to the existence of $w_1$ and $w_2$ letting $D(w_1)G - CD(w_2) = 0$. First, if $E(G, C) = 0$, by definition, there exists a diagonal matrix $D^\star$ such that $E_s(G, CD^\star) = 0$. Note that normalization to simplex is done by a left diagonal matrix multiplication, e.g., for $G$, the normalized growth rates can be given by $D_G G$ where the $i$th diagonal element of $D_G$ is the sum of the $i$th row of $G$. The fact $E_s(G, CD^\star) = 0$ suggests that $D_G G = D_C CD^\star$ where $D_C$ normalizes rows of $CD^\star$ to the simplex, and leads to the existence of $w_1$ and $w_2$: $D(w_1) = D_C^{-1} D_G$ and $D(w_2) = D^\star$. On the contrary, if the positive $w_1$ and $w_2$ making $D(w_1)G - CD(w_2) = 0$ exist, we can directly have $E_s(G, CD(w_2)) = 0$ and then $E(G, C) = 0$ since $E(G, C) \le E_s(G, CD(w_2)) = 0$ and $E(G, C) \ge 0$ by definition. The encroachment, $E(G, C)$, therefore also has strong relations with global stability, providing a sufficient condition of global stability ($E(G, C) = 0$).

However, it is not well-understood why in simulation results (e.g., Fig. 4), global stability begins to be lost near $E(G, C) = 0.5$. Also, for other resource supply method, it seems difficult to prove that $E(G, C) = 0$ must lead to global stability which is likely correct based on simulation results. In a word, global stability is still mysterious and deserves future studies.

#### S-II. NUMERICAL METHODS

This section will introduce how numerical experiments were conducted. We mainly use Julia to run simulations and Python to analyze the results. The specific codes can be found online at github.com/liuyz0/RCM-stability. Usually, there are several modules in each simulation process:

- Determining system size: We need to first specify the number of species and resources at the given fixed point, i.e., $N_S$ and $N_R$.

- Sampling: Given $N_S$ and $N_R$, we then sample different consumption rates $C$, as well as growth rates $G$. The fixed point, embodied by equilibrium species abundances $S^*$ and equilibrium resource concentrations $R^*$ will also be sampled. Based on different resource supply methods, some other parameters will need to be sampled. The rest parameters can be solved based on previously sampled ones and the condition that time derivatives are all zero at the given fixed point.

- Simulating: With the sampled parameters, ordinary differential equations, Eq. (S1), are simulated with various initial conditions. Key results like the number of surviving species, whether the community is fluctuating or not, and whether the community has alternative stable states, will be recorded.

- Analyzing data: A part of the data, for instance, the Jacobian, Eq. (S13), the characteristic matrix $GC^\top$, and the encroachment $E(G,C)$, can be directly calculated and analyzed at once after sampling. Other results, like fraction of survival, are obtained during dynamics simulations. Each community can produce various results and data from millions of communities will be averaged or compressed according to certain perspectives of our theory, yielding the organized results in the figures.

Next, we explain in detail about these steps one by one.

##### A. Sampling methods

After given the system size, we first sample the two $N_S \times N_R$ matrices $C$ and $G$. In the paper, there are two sampling ways, one is with strict metabolic trade-off ensuring uniform equilibrium resource concentrations (special sampling), and the other samples everything including the fixed point randomly (general sampling). The metabolic trade-off requires growth vectors $G_i$ to be on the simplex. To obtain most diverse growth vectors, we sample $G_i$ uniformly from the simplex. Since a $G_i$ is a $N_R$ vector, we first sample $N_R - 1$ i.i.d. variables $X_\alpha$ uniformly from $[0,1]$ (i.e., $\mathcal{U}[0,1]$). Sorting these variables from small to large and getting $X_\alpha^\uparrow$ (i.e., $X_1^\uparrow \le X_2^\uparrow \le \cdots \le X_{N_R-1}^\uparrow$, and $\{X_\alpha^\uparrow\}_{\alpha=1}^{N_R-1} = \{X_\alpha\}_{\alpha=1}^{N_R-1}$), we can obtain a $N_R$ vector $(X_1^\uparrow - 0, X_2^\uparrow - X_1^\uparrow, ..., X_{N_R-1}^\uparrow - X_{N_R-2}^\uparrow, 1 - X_{N_R-1}^\uparrow)$, which can be shown to be uniformly distributed on the simplex. Independently sampling $N_S$ such vectors can result in the full matrix $G$. The general sampling method is simpler, which samples each element $G_{i\alpha}$ from $\mathcal{U}[0,1]$. In sampling $C$, we first use the same sampling as $G$ to get another $N_S \times N_R$ matrix $C_0$, and let

$$C = \rho G + \sqrt{1-\rho^2}C_0. \tag{S36}$$

Clearly, by varying $\rho$ from 1 to 0, we can realize different configurations from $C = G$ to $C$ independent of $G$. In the general sampling scheme, we further rescale $C$ by sampling a random positive diagonal matrix (elements from $\mathcal{U}[0.01,1]$) and multiplying $C$ from the right side.

In the general sampling scheme, the elements of $S^*$ and $R^*$ are all independently drawn from $\mathcal{U}[0.01,1]$. The mortality $\delta$ can be solved readily from $\mathrm{d}S/\mathrm{d}t = 0$ as $\delta = GR^*$. With metabolic trade-off, given the mortality is universally 1, we have $R_\alpha^* = 1$, $\forall\alpha$, while $S^*$ are still drawn from $\mathcal{U}[0.01,1]$.

For constant resource supply, $h_\alpha(R_\alpha) = \gamma_\alpha$, the supply constants, $\gamma_\alpha$ can be directly solved from $\mathrm{d}R/\mathrm{d}t = 0$ given $R^*$, $S^*$, and $C$. For linear supply, $h_\alpha(R_\alpha) = l_\alpha(\kappa_\alpha - R_\alpha)$, we sample $l_\alpha$ i.i.d. from $\mathcal{U}[0.1,1]$ and solve $\kappa_\alpha$. And for logistic supply, $h_\alpha(R_\alpha) = g_\alpha R_\alpha(K_\alpha - R_\alpha)$, $K_\alpha$ will be solved out after sampling $g_\alpha$ i.i.d. from $\mathcal{U}[0.1,1]$.

##### B. Simulation methods

After obtaining all the parameters needed, we can begin to simulate dynamics. In Julia, we use the `Differential Equations` package and its solvers. Most times, the dynamics are non-stiff, and we tried solvers like `Tsit5` (Tsitouras 5/4 Runge-Kutta method), `Vern7` (Verner's "Most Efficient" 7/6 Runge-Kutta method), and `VCABM3` (the 3rd order Adams-Moulton method), which have no obvious differences in results. However, for logistic resource supply, when $C$ is almost independent of $G$, the dynamics can be stiff and may require stiff solvers like `Rodas4` (4th order A-stable stiffly stable Rosenbrock method with a stiff-aware 3rd order interpolant) and `TRBDF2` (a second order A-B-L-S-stable one-step ESDIRK method). A `callback` function is needed to force the solutions to be non-negative, preventing non-physical results and maintaining numerical stability.

Before simulating the dynamics, we also need to determine initial conditions. For each community, there are two ways of setting the initial values depending on the purpose: one is perturbing the given fixed point (initial variable values of are sampled uniformly in $0.95 \sim 1.05$ of the corresponding equilibrium values, respectively); the other is to randomly sample them (initial values are sampled uniformly in $0.005 \sim 1.995$ of the corresponding equilibrium values, respectively).

Specific values of different cutoffs and the simulation procedures are introduced as follows. Most communities with $G \approx C$ or $C$ nearly independent of $G$ can converge within time `1e+3`, while those at the edge of being stable/unstable need much longer time (typically $\sim$ `1e+5`) to show evidences of convergence/divergence. To best identify fluctuations while reduce simulation time cost, we check the convergence every $\Delta t =$ `5e+3`. We add a small constant dispersal $D =$ `1e-7` to species during simulation. On the one hand, we can ensure positiveness of abundances. On the other hand, we can set a criterion for extinction, i.e., abundance lower than `1e-5` will be regarded as extinction, since the contribution of dispersal to abundance alone during each period can reach $\Delta t D \sim$ `1e-4`. At the end of each $\Delta t$ period, we find the survival species first, and check the average (over surviving species) relative difference between species abundances at the end and the mean values (over the last `1e+3` time). By manually, checking, the average relative difference for a converging community is typically smaller than `1e-3` while that for a fluctuating community is much larger, i.e., of the order `1e-1` $\sim$ `1e+0`. Therefore, we set the critical value as `1e-3` to distinguish stable and fluctuating communities. If a community is judged as being stable, we will end the simulation. Otherwise, we will continue a new $\Delta t$ period of simulation. The upper bound of the number of periods is 15. If a community does not converge after $15\Delta t$, we will regard it as a fluctuating one. For fluctuation case, we can further use Julia package `DynamicalSystems` and the function `lyapunov` to identify the oscillatory attractor found is chaotic or not. With these procedures, we can obtain survival fraction, whether a system is fluctuating, and the type of fluctuation, given the parameters and one set of initial conditions.

Since we want to distinguish alternative stable states and globally stable states, we need to test different initial conditions for a fixed set of parameters. Among all results obtained from different initial conditions, those go to stable states will be identified as globally stable as long as (i): there is no fluctuation starting from any initialization; and (ii): the stable states are the same fixed point. Otherwise, we will call these stable states as alternative stable states. We use principal component analysis (PCA) to tell whether different results in the space belongs to the same fixed point. One converging simulation case yields final species abundances $S$ and resource concentrations $R$, which forms a characteristic vector $(S; R)$. For `num_init` different initial conditions, we can have a `num_init` by `num_init` matrix with the $ij$ element being $\cos\theta_{ij}$ where $\theta_{ij}$ is the angle between the $i$th and $j$th characteristic vectors. PCA calculates the eigenvalues of such a matrix. We count the number of eigenvalues larger than 5% of the largest eigenvalues, which is regarded as the number of different components or stable states. Global stability will be concluded if this number is 1. With all these methods, we now are able to distinguish global stability, multistability, fluctuation, etc. for each simulation case. With proper classification of the communities, like value of encroachment, $E(G, C)$, we can calculate the fractions of different types of dynamics inside each class, as well as mean survival fraction.

##### C. Data analysis methods

Once the parameters are sampled, the Jacobian, the matrix $GC^\top$, and encroachment $E_s(G, V)$ or $E(G, C)$ can be calculated directly. Most quantities can be directly and simply calculated by definition. The calculation of $E(G, C)$ is a little different since optimization is involved. We used PyTorch in python or Flux in Julia to write self-defined layer to output $CD$ with $C$ as the input and $D$ the parameters to optimize. The loss function is given by $E_s(G, CD)$. We tried different solvers like Adam and stochastic gradient descent (SGD) which yield the same results. The learning rate is set as `lr` $=$ `0.1` and the number of iterations is `1000`. These settings have been tested to guarantee convergence for all optimization processes.

When we have all the results needed for each simulation case, we do statistics based on different purposes. For example, if we want to see how certain value changes with respect to encroachment, we first divide all the communities into many groups with each group having similar $E(G, C)$. And within each group of communities, we average the value of interest across communities. At last, we obtain a relation between the mean of this quantity and the encroachment.

#### S-III. FIGURE DETAILS AND SUPPLEMENTARY RESULTS

In this section, we present here the detailed settings of numerical experiments shown in figures. Table S1 briefly summarize the settings for each figure which will be discussed one by one. Supplementary results supporting the arguments in the main text and verifying the validity of our method for more cases will also be discussed along the way.

| Settings / Figures | System size | Sampling method | Initial condition | Sample range |
|---|---|---|---|---|
| Fig. 1, C and D | $N_R = 64,\ N_S = 32$ | General | Perturbing | $\rho = 0.4$ or 1.0 |
| Fig. 2, D and E | $N_R = 32,\ N_S = 16$ | Special | | Four different mean niche ranges (by restricting sampling on simplex to be concentrate); for each mean niche range, try 512 $\rho$ values ranging from 0 to 1; for each $\rho$ value, sample one set of $G$, $C$, and other parameters. |
| Fig. 3, A and D | $N_R = 32,\ N_S = 16$ | General | | Four different equilibrium resource concentration distributions; for each distribution, try 512 $\rho$ values ranging from 0 to 1; for each $\rho$ value, sample one set of $G$, $C$, $S^*$, and $R^*$. |
| Fig. 3E | $N_R = 32,\ N_S = 16$ | General | Perturbing | 144 different $\rho$ from 0 to 1; for each $\rho$, sample one pair of $G$ and $C$; for each pair of $G$ and $C$, sample 10 different fixed points; for each case, try 10 different initial conditions in simulation. |
| Fig. 3E internal | $N_R = 32,\ N_S \in 4:48$ | General | | For each $N_S$, try 144 different $\rho$ from 0 to 1; for each $\rho$, sample one pair of $G$ and $C$; for each pair of $G$ and $C$, sample 10 different $S^*$ and $R^*$. |
| Fig. 4 | $N_R = 32,\ N_S = 16$ | General | Non-perturbing | 1440 different $\rho$ from 0 to 1; for each $\rho$, sample 10 different sets of $G$, $C$, $S^*$, and $R^*$; for each case, try 50 different initial conditions in simulation. |
| Fig. 5 | $N_R = 32,\ N_S \in 4:48$ | General | Non-perturbing | For each $N_S$, try 144 different $\rho$ from 0 to 1; for each $\rho$, sample one pair of $G$ and $C$; for each pair of $G$ and $C$, sample 10 different fixed points; for each case, try 10 different initial conditions in simulation. |

TABLE S1. A summary of numerical experiment settings in different main text figures. Definitions of the terms in the table can be found in Sec. S-II A and S-II B.

##### A. Figure 1

The simulation in Fig. 1 is for demonstration. We set up a community with $N_R = 64$, $N_S = 16$, and give the fix point at $S_i^* = 1$ and $R_\alpha^* = 1$. The matrices $G$ and $C$ are sampled in the general way, and the initial condition is obtained by perturbing the given fixed point (see Sec. S-II A). When the parameter $\rho$ in Eq. (S36) controlling correlation between $G$ and $C$ is 1.0, we have Fig. 1C. Whereas, when $\rho = 0.4$, we can obtain dynamics shown in Fig. 1D.

##### B. Figure 2

We did not simulate the dynamics in Fig. 2 but only sampled the communities and examined the stability via the Jacobian, and different criteria. The communities are still of size $N_R = 32$, $N_S = 16$. Samples of $G$ are restricted on the simplex to satisfy the metabolic trade-off (see Sec. S-II A). Since we know the center of the simplex is $(1, 1, ..., 1)/N_R$, after uniformly sampling the vectors $G_i$ on the simplex, we can use $G_i = \texttt{comp}(G_i - (1, 1, ..., 1)/N_R) + (1, 1, ..., 1)/N_R$, where $0 \le \texttt{comp} \le 1$, to make $G_i$ uniformly on the small interior of the simplex and tune mean niche ranges. We chose four values of `comp`: 1.0, 0.8, 0.6, and 0.4. For each `comp` value, we sampled 512 different $\rho$ (Eq. (S36)) values. And for each $\rho$, we sample one community with the special way (with metabolic trade-off and uniform equilibrium resource concentrations). Calculating the simple encroachment $E_s(G, V)$ and the interaction measure (Eq. (S28)) for each sampled community and doing statics, we obtained Fig. 2 D and E.

We demonstrate more cases supplementary to Fig. 2 B and C. In the main text, we explain instability based on flipping of impact vectors for two species with two resources. In high dimensional cases, the configuration of mutual flipping between two species can still happen but it is not the only configuration leading to instability. As shown in the left of Fig. S3 for three species, species $i$ can consume in species $j$'s niche while species $j$ does not consume in species $i$'s niche. Therefore, mutual flipping is not present. However, the system can be unstable or even have persistent fluctuation since we have a paper-rock-scissor type interaction between species (species 1 suppresses 2, species 2 suppresses species 3, and species 3 suppresses species 1). We also emphasize that encroachment greater than 1 is a necessary condition for instability as it enable species to consume in others' niches but does not ensure this. As shown in the right of Fig. S3 for two species and three resources, when encroachment is greater than 1, there may not be flipping between impact vectors. In conclusion, we make the arguments mentioned in the main text more concrete here that there are more instability configurations in high dimensional cases and encroachment greater than 1 is a necessary condition for instability.

##### C. Figure 3

In Fig. 3, we fixed system size as size $N_R = 32$, $N_S = 16$ and used the general sampling method for $G$, $C$, and the given fixed points (see Sec. S-II A). For Fig. 3 A and D, the equilibrium resource concentrations are not totally random though. We chose four different resource concentrations, $R^*$, with different evenness. For each resource concentration vector $R^*$, we sample 512 communities (each community was assigned with one $\rho$ (Eq. (S36)) value) whose growth rates, $G$, consumption rates $C$, and equilibrium species abundances $S^*$ were randomly chosen. Left parameters of each community were solved from the given fixed point. Calculating the real Jacobian and encroachment defined for each community led to Fig. 3 A and D, where simple encroachment fails to characterize instability for non-uniform resource concentrations, while the newly defined encroachment can.

In Fig. 3E, with the same system size, we tried 144 different $\rho$ values (Eq. (S36)). For each $\rho$, one pair of $C$ matrix and $G$ matrix was sampled. To emphasize the point that stability robustly depends only on $G$ and $C$, for each pair of $C$ and $G$, we sampled 10 different communities giving distinct fixed points. We initialized the simulation by perturbing the given fixed point (see Sec. S-II B). And 10 different initial conditions were tried for each community. Fraction of survival was recorded for each community, and within each small interval (having length 0.1) of encroachment $E(G, C)$, the quantity was averaged over communities inside the interval yielding Fig. 3E.

The inner figure of Fig. 3E use the same sampling method as Fig. 3E while also tried $N_S$ from 4 to 48. The real Jacobian and $GC^\top$ were calculated and the fraction of unstable modes defined as Eq. (S17) of each matrix was recorded. As a result shown in the internal figure, on average, $GC^\top$ has the same fraction of unstable modes as the Jacobian.

We test our instability criteria for different resource supply methods and derive the semi-empirical law for survival fraction here. Supplementary to the main text, we also considered linear resource supply and logistic resource supply. Given resource supply method and system size $N_R = 32$, $N_S = 16$, we tried 144 different $\rho$ values (Eq. (S36)) in sampling $G$ and $C$. Given a $\rho$, one pair of $C$ matrix and $G$ matrix is sampled. For each pair of $C$ and $G$, we sampled 10 different communities giving distinct fixed points. We initialized the simulation by perturbing the given fixed point (see Sec. S-II B). And 10 different initial conditions were tried for each sampled community. The fraction of unstable modes of the Jacobian for constant resource supply and linear resource supply models are given by Eq. (S17). However, for logistic resource supply model, we re-define the fraction of unstable modes as

$$\frac{\#\text{ of eigenvalues with positive real parts}}{N_S + N_R}, \tag{S37}$$

Using the criterion determining whether a community is stable or not based on the Jacobian, we examine the fraction of unstable communities given a specific encroachment value. The result shows that encroachment $E(G, C)$ work well for the resource-consumer model regardless of resource supply methods (Fig. S1A). Encroachment value keeps to work in predicting survival fraction for different resource supply methods (Fig. S1B). The survival fraction begins to drop when encroachment is near 1, and then decreases linearly as

$$\text{Survival fraction} = \frac{3}{2} - \frac{1}{2}E(G, C),\ \ E(G, C) \geq 1, \tag{S38}$$

which is the line drawn in Fig. S1B. To understand this linear relation, we first connect fraction of survival to that of unstable modes, and then connect encroachment to the fraction of unstable modes of the Jacobian. The empirical results show that on average, one unstable direction leads to the extinction of two species in the dynamics ($y = 1 - 2x$ in Fig. S1C where $x$ is the fraction of unstable modes and $y$ the survival fraction). This also justifies the need of the concept statistical stability based on the fraction of unstable modes, since it is the fraction of unstable modes making the complex system different for observers (e.g., observables like survival fraction can have difference). Next, we find that encroachment has a clear relation with the fraction of unstable modes of the Jacobian as is argued in

Sec. S-I D. The relation is well approximated by $y = (x-1)/4$ (solid line in Fig. S1D) where $x$ denotes encroachment and $y$ is the fraction of unstable modes. Combining relationships between unstable mode fraction and encroachment and that between survival fraction and unstable mode fraction, we obtain the formula Eq. (S38). In summary, the supplementary results support validity of encroachment analysis for all three resource supply methods, justify the importance of the concept, statistical stability, which is defined based on unstable mode fraction, and reveal that encroachment well captures the fraction of unstable modes–the average degree of instability for the community.

We also directly check if encroachment can well reflect stability through the lens of highly diverse finial communities. In practice, we evaluate the fraction of communities going to high survival fraction (= 1.0) stable states as the fraction of being stable, which does begin to decrease near $E(G, C) = 1$ (Fig. S1E).

Finally, we check the validity of the characteristic matrix $GC^\top$ in identifying stability for all three resource supply methods. Results show that the characteristic matrix $GC^\top$ has the same fraction of unstable modes as the Jacobian $J$ on average (Fig. S1F), suggesting that $GC^\top$ continues to work regardless of what kind of resources are supplied.

##### D. Figure 4

We studied dynamics beyond perturbing the fixed point in Fig. 4. Given the system size $N_R = 32$, $N_S = 16$, we used 1440 different $\rho$ to sample. For each $\rho$ (Eq. (S36)), 10 communities were obtained via the general sampling way (see Sec. S-II A). For each community, we simulated the dynamics starting from 50 different randomly chosen initial conditions not perturbing the fixed point but distributed in the space (see Sec. S-II B). Different fractions were recorded based on the procedure in Sec. S-II B. The mean value of certain quantity with respect to encroachment was obtained by averaging that quantity over communities with the same encroachment value (in practice, communities within the same encroachment interval of length 0.1).

We present here dynamics of resource-consumer models with different kinds of resource supply methods (i.e., constant, linear, and logistic supply). The system size, sampling method, initial conditions, and simulation procedures are the same as those for Fig. 4 but with different resource supply methods. Fraction of survival species and those of various dynamical behaviors are evaluated and compared (Fig. S4, A-H). Different resource supply methods have similar instability transition threshold, i.e., $E(G, C) = 1$, while communities with logistic resource supply tend to lose stability a little earlier (see fraction of survival of stable states and survival fraction of fluctuating state in Fig. S4, A and B, respectively). However, for dynamical behaviors, only constant and linear resource supply methods are similar, which have huge differences comparing to the logistic resource supply. When $E(G, C) > 1$, the fraction of fluctuation/alternative stable states of communities with logistic resource supply can be much larger/smaller than those of communities with other resource supply methods (Fig. S4E/F). When looking into the details of fluctuating communities, classifying fluctuations into chaos or limit cycles, we find that the majority of fluctuations with logistic resource supply is chaotic while for the other two resource supply methods, the majority is limit cycles (see Fig. S4, G and H for chaos and limit cycles, respectively). In conclusion, the dynamic simulations also support encroachment to be a good pointer for instability regardless of resource supply methods, while the specific dynamical behaviors may be affected by what kind of resources are supplied and need future works to explain.

##### E. Figure 5

To include the effect of species-resource ratio, $N_S/N_R$, we tested $N_S$ from 4 to 48 and $N_R = 32$ in Fig. 5. For each $N_S$, 144 different $\rho$ (Eq. (S36)) were used. For each $\rho$, we sampled one pair of $G$ and $C$ via the general sampling way (see Sec. S-II A). And for each pair of $G$ and $C$, we sampled 10 different communities by assigning distinct given fixed points and solving the rest parameters. For each community, 10 different initial conditions were tried in the simulation which were not perturbed from the fixed points (see Sec. S-II B). For the two dimensional phase diagrams, the color at each pixel representing certain mean value were calculated by averaging the quantity over communities within that pixel (i.e., communities having the same $N_S/N_R$ and encroachment $E(G, C)$). Each pixel corresponds to one $N_S/N_R$ value and has a length 0.1 in the axis of encroachment.

We derive the semi-empirical scaling law that predicts the exact stability boundary considering $N_S/N_R$ as follows. When encroachment $E(G, C) > 1$, it is possible for instability configurations (e.g., mutual flipping of impact vectors) to happen, while it is not necessary for such configuration to happen (see Fig. S3 Right). Without loss of generality, we can assume there is a probability $P_u(E)$ for a pair of neighboring species to have flipping. The probability $P_u(E)$ begin to be non-zero when encroachment $E = 1$ and will increase thereafter with increasing $E$. Intuitively, $P_u(E)$ can be understood as the ratio of niche range intersection area to the area of niche, and thus should also depends on $N_R$, the geometric dimension of the niche ranges. The number of pairs of neighboring species is smaller than $N_S$ but greater than $N_S/2$. Therefore, the probability of at least one pair can have flipping is given by

$$1 - (1 - P_u(E))^{\Omega(N_S)}, \tag{S39}$$

where $\Omega(N_S)$ denotes some function scaling as $N_S$ (up to some constant). Assume when the probability $1 - (1 - P_u(E))^{\Omega(N_S)}$ is large enough, i.e., larger than $1 - \epsilon$ for sufficiently small $\epsilon$, we can be almost sure to observe instability, then the encroachment sufficiently leading to instability should satisfy

$$P_u(E) \geq 1 - \epsilon^{\frac{1}{\Omega(N_S)}}. \tag{S40}$$

Since $P_u(E)$ is always small even after $E$ changes from 1 to 2, we use a linear function (or only take the first order Taylor expansion) to approximate the small change of $P_u(E)$: $P_u(E) \propto E - 1$. Therefore, we obtain the critical encroachment as

$$E = 1 + a(1 - \epsilon^{\frac{1}{\Omega(N_S)}}), \tag{S41}$$

for some constant $a$. Combining with the understanding that the critical encroachment value that sufficiently leads to instability should depends on $N_S/N_R$, we refine the above equation by adding $N_R$ based on how it depends on $N_S$ to get the scaling of critical encroachment:

$$E = 1 + a(1 - b^{\frac{N_R}{N_S}}), \tag{S42}$$

where $b$ should be some constant. When $N_S/N_R \ll 1$, the critical encroachment is observed to be  1.7 (Fig. 5), such that we set $a = 0.7$. The curve given by Eq. (S42) is not very sensitive to the choice of $b$ ($b$ from 0.5 to 0.8 gives similar results). We choose $b \sim 0.65$ to get the boundary drawn in Fig. 5. Notably, we should also consider instability configurations other than mutual flipping (e.g., Fig. S3 Left). But this will not change how the formula scales with $N_S$ as the number of other configurations also scale up proportionally to $N_S$. Also, since we focus on statistical stability, to be rigorous, we should consider the probability that a fraction of species are involved in instability configurations rather than the probability of having any instability configuration. However, as long as the fraction is small, we believe the basic scaling behavior is still captured by Eq. (S42), and thus it can agree well with the boundaries of losing species or having fluctuations in Fig. 5.

We examine the phase diagrams for different resource supply methods in Fig. S5 where the simulation begins by perturbing the given fixed points. For initial conditions not perturbing the fixed points, the simulation results for different resource supply methods are presented in Fig. S6. Fraction of alternative stable states when $N_S/N_R > 1$ increases when we randomly sample the initial conditions in a wide range. We study the effect of $N_R$ on numerical results in Fig. S7, which suggests that $N_R = 32$ is large enough to show the features of complex systems in a sense the boundaries are smooth, clear, and has small difference comparing to greater $N_R$.

---


[S1] R. M. Arthur, Proceedings of the National Academy of Sciences **64**, 1369 (1969).
[S2] D. Tilman, *Resource Competition and Community Structure* (Princeton University Press, 1982).
[S3] Y. Ahmadian, F. Fumarola, and K. D. Miller, Physical Review E **91**, 012820 (2015).
[S4] L. Stone, Scientific Reports **8**, 8246 (2018).

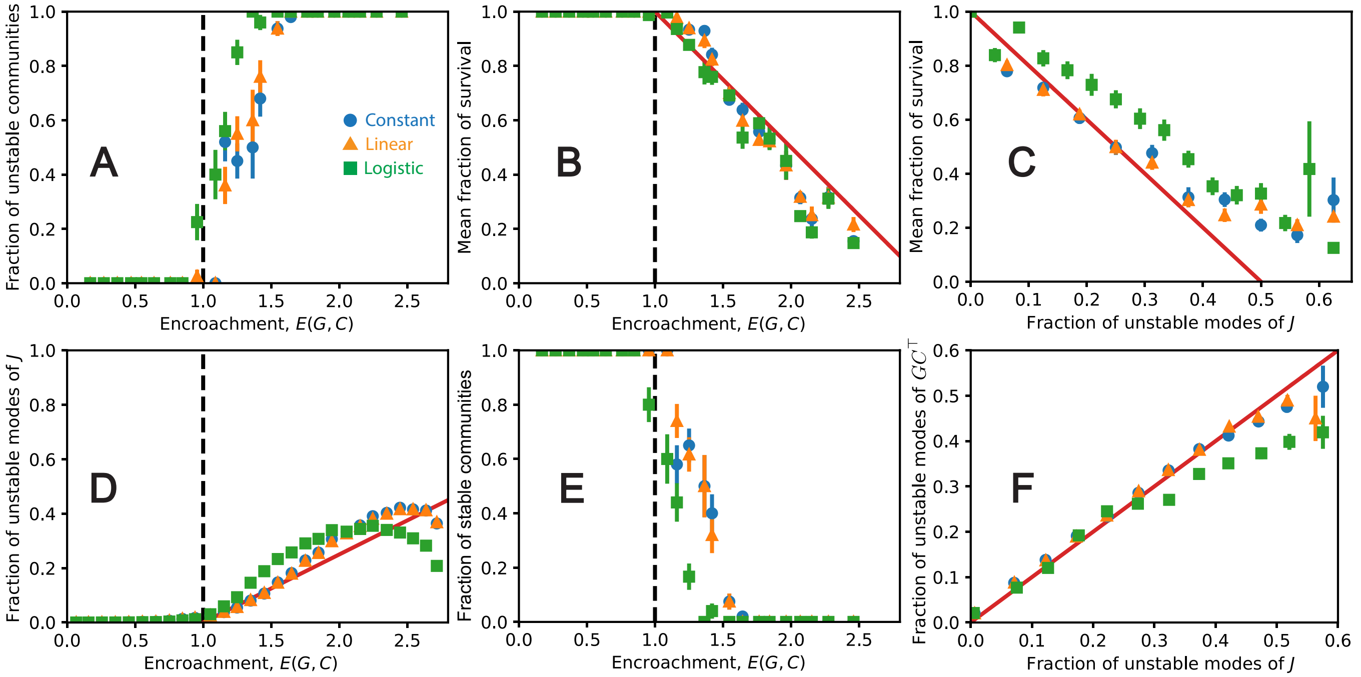


FIG. S1. Supplementary results for stability show the encroachment analysis works for all three resource supply methods and provide a semi-empirical analysis of species survival fraction: (A) Different methods of resource supply (● for constant supply, ▲ for linear supply, and ■ for logistic supply) do not change the fact that instability transition happens at $E(G, C) = 1$. (B) Fraction of survival begins to drop near $E(G, C) = 1$ and then show approximately a linear relation with encroachment $E(G, C)$. (C) For all three resource supply methods, it is roughly true that every unstable direction leads to the extinction of two species (the slope of the solid line is $-2$), such that the fraction of survival has a linear relation with the unstable mode fraction of the Jacobian $J$. (D) Encroachment as an indicator of instability naturally has a relation with the fraction of unstable modes of the Jacobian (the solid line captures initial change of unstable mode fraction with respect to encroachment, which intersect x-axis at $E(G, C) = 1$ and has a slope 0.25). (E) We evaluate the fraction of fully coexisting stable states which sharply vanish when encroachment reaches 1. (F) For all three resource supply methods, the characteristic matrix $GC^\top$ has the same fraction of unstable modes as the Jacobian $J$ on average.

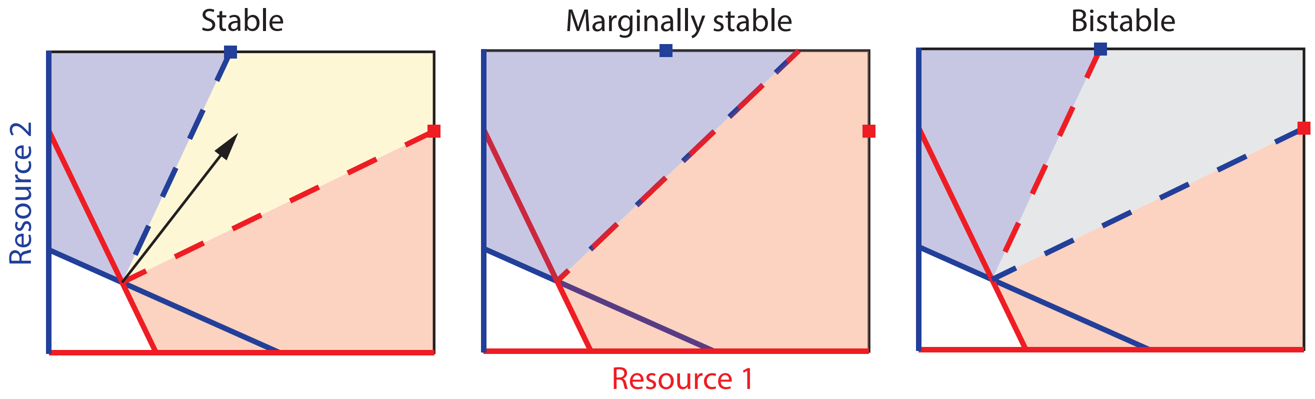


FIG. S2. Tilman's graph shows feasibility and stability via geometries in resource space for simple communities with two species and two resources. Axes represent resource concentrations and solid lines in the figures are ZNGIs of species. In the left patch, the ZNGIs intersect and the supply vector (black arrow) is between the dashed lines (parallel to impact vectors), such that the system has a feasible equilibrium. Besides, impact vectors are more aligned with normal vectors of ZNGIs, so the system is stable. In the middle patch, the impact vectors begin to flip, which leads to marginal stability. Impact vectors in the right patch have flipped, such that species tend to consume resources the other one grow on and the coexisting feasible point is unstable.

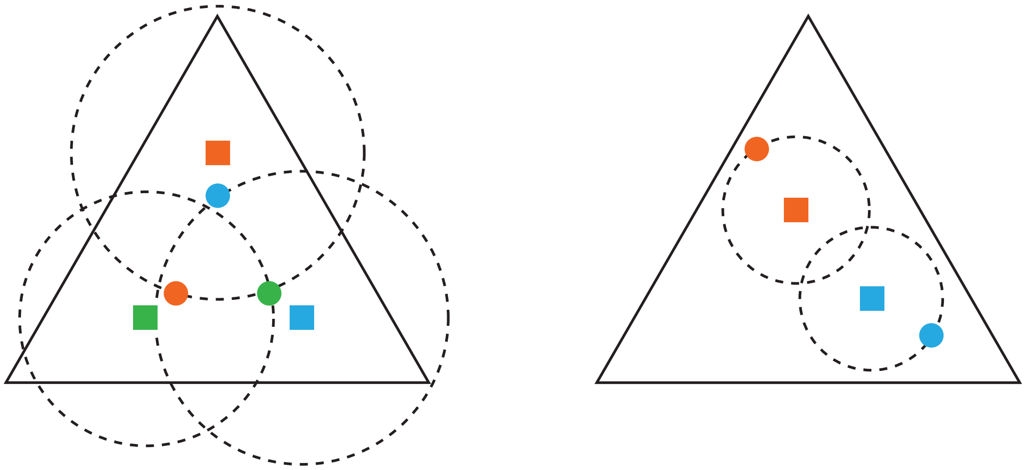

FIG. S3. High dimensional cases can have more configurations leading to instability and the limitations of encroachment analysis. Squares/dots in the figure represent growth/impact vectors normalized on simplex. (Left) Unlike the cases with two species and two resources, in high dimensional cases, like with three species and three resources, there can be more configurations leading to instability rather than mutual flipping of impact vectors. In the figure, we demonstrate such an example where species $i$ consume species $j$'s in niche for $i = 1, 2, 3$ and $j = 2, 3, 1$, respectively, which may lead to fluctuations. (Right) Encroachment $E(G, C) > 1$ is technically only a necessary condition for instability. As shown in the figure, when $E(G, C) > 1$ for two species, there might not be flipping of impact vectors and the system can still be stable.

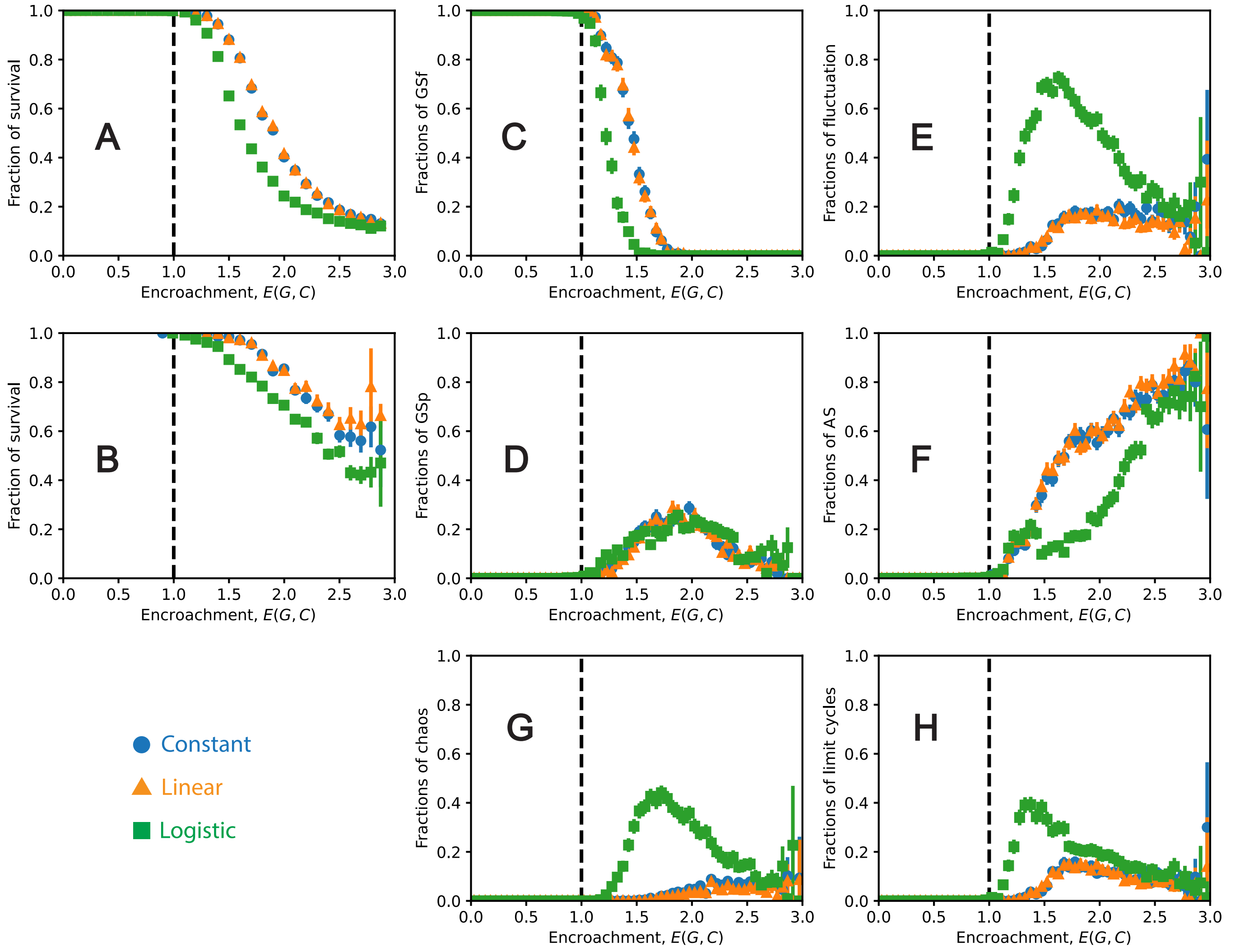

FIG. S4. Results of dynamics of all resource supply methods and a detailed exploration of alternative stable states. In all the simulations, we initialize the variables randomly. In (A)-(H), we use • to denote constant supply, ▲ for linear supply, and ■ for logistic supply. (A) Fraction of survival species of stable states begin to drop when encroachment reaches 1. (B) Fraction of survival species of fluctuation states begin to drop when encroachment reaches 1. (C) Fractions of community going to globally stable and fully coexisting states decreases near $E(G, C) = 1$. (D) Fractions of community going to globally stable but partly coexisting states increases near $E(G, C) = 1$ for all resource supply methods. (E) Fraction of communities going to fluctuating states begin to increase near $E(G, C) = 1$. (F) Fraction of communities going to alternative stable states begin to increase near $E(G, C) = 1$. (G) Investigating fluctuating communities in detail, we show the fraction of communities going to chaotic fluctuation among all communities. (H) The fraction of communities going to limit cycles among all communities are also shown.

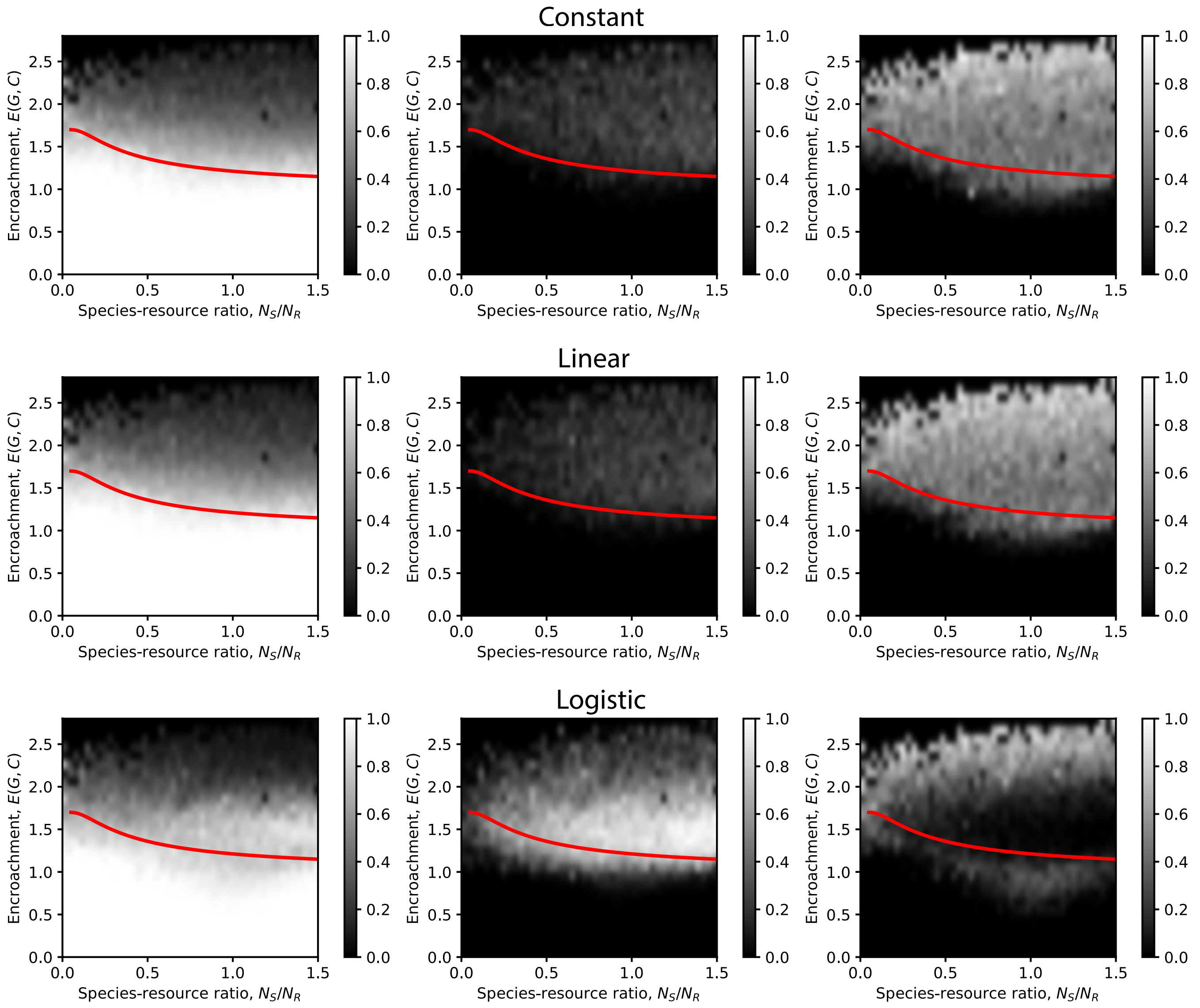


FIG. S5. Phase diagrams of dynamical behaviors (columns: first column is survival fraction, the second is fluctuation fraction, and the last is fraction of alternative stable stats) of communities with sizes $N_R = 32$, $N_S$ from 4 to 48, and different resource supply methods (rows). The initial condition is given by perturbing the given fixed point in each simulation case (see Sec. S-II B). The line inside is the scaling law given by Eq. (S42).

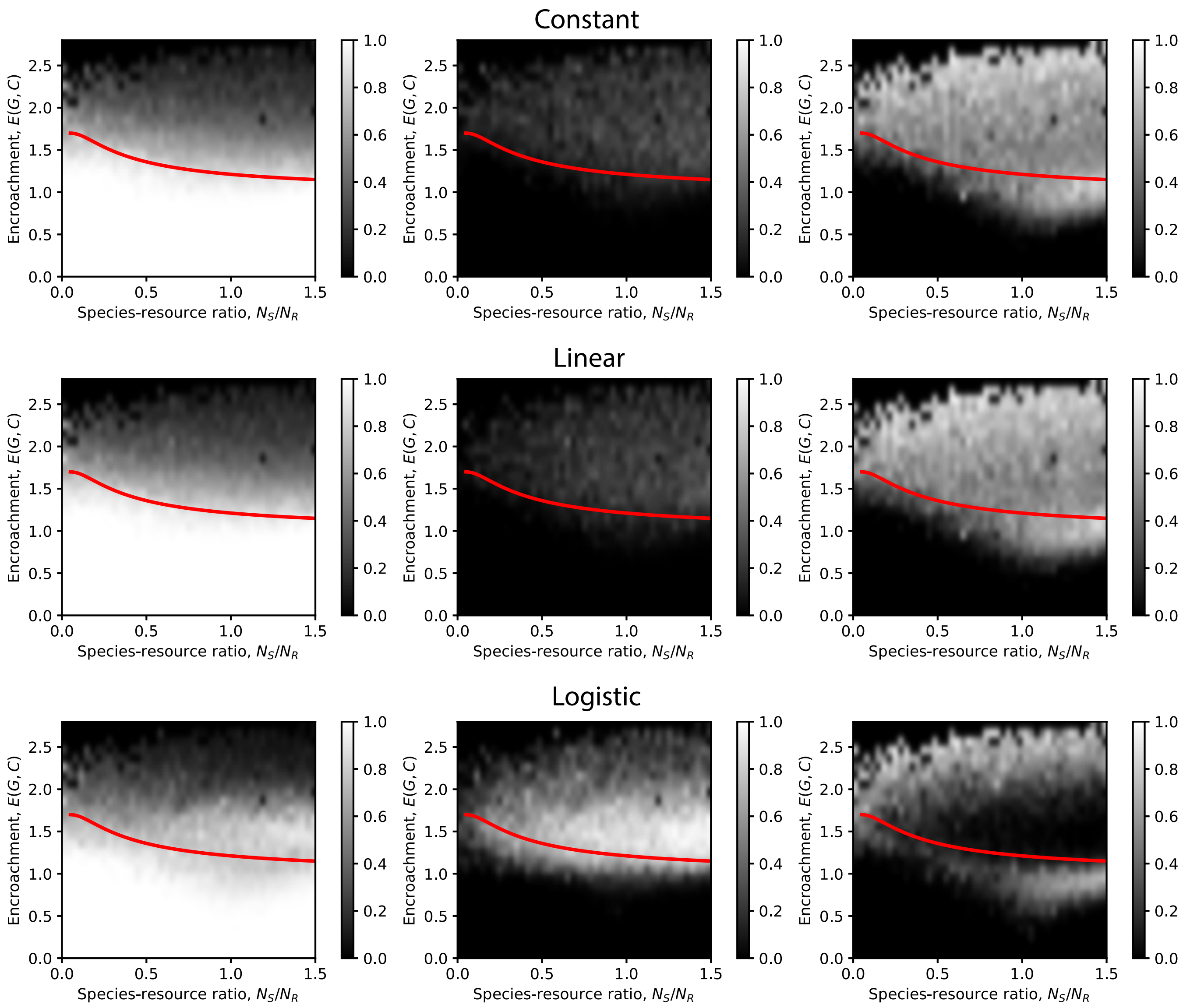


FIG. S6. Phase diagrams of dynamical behaviors (columns: first column is survival fraction, the second is fluctuation fraction, and the last is fraction of alternative stable stats) of communities with sizes $N_R = 32$, $N_S$ from 4 to 48, and different resource supply methods (rows). The initial condition is sampled in a wide range (non-perturbing) in each simulation case (see Sec. S-II B). The line is the scaling law given by Eq. (S42).

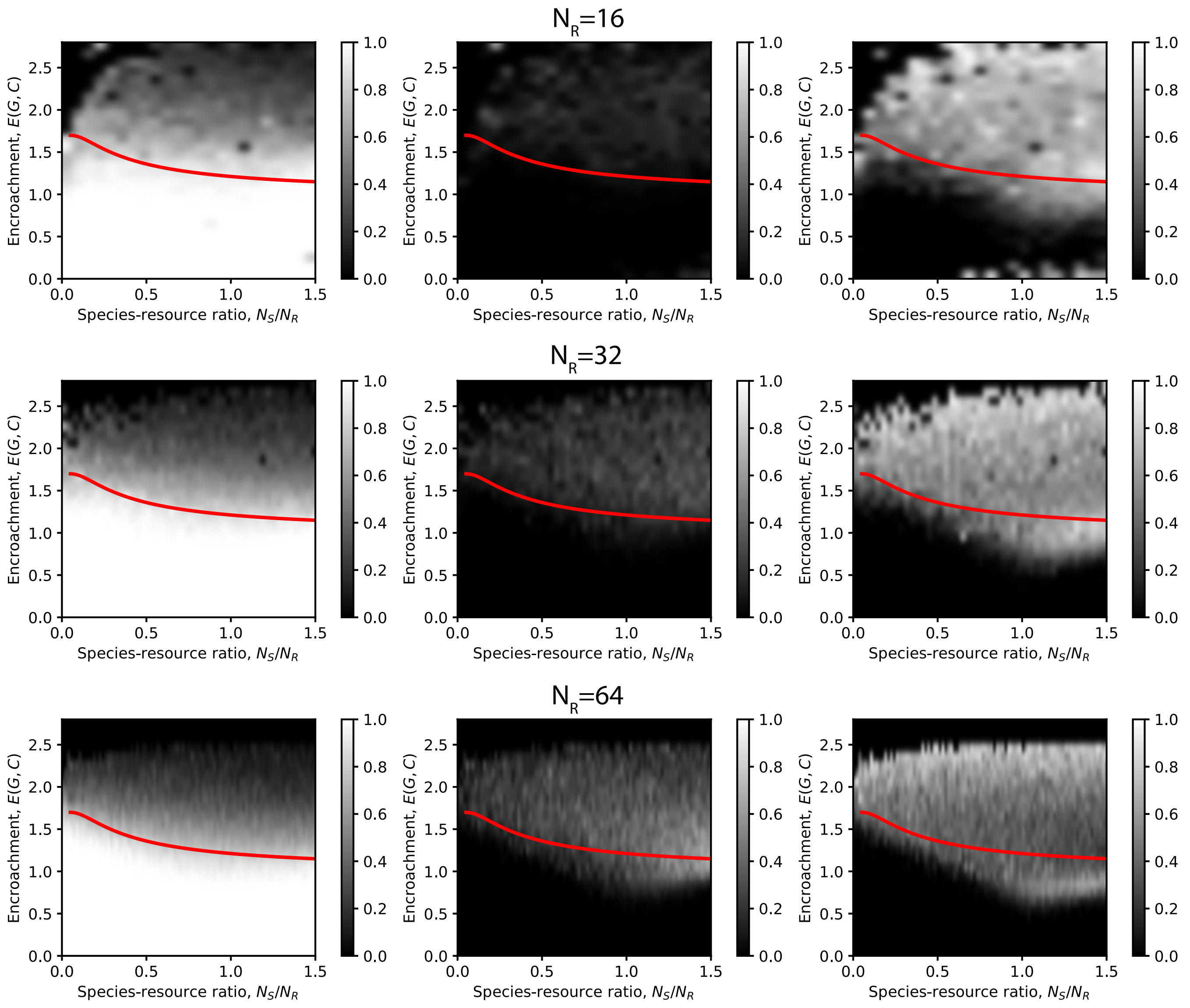


FIG. S7. Phase diagrams of dynamical behaviors (columns: first column is survival fraction, the second is fluctuation fraction, and the last is fraction of alternative stable stats) of communities with different system sizes (rows) and the constant resource supply method. The initial condition is sampled in a wide range (non-perturbing) in each simulation case (see Sec. S-II B). The line is the scaling law given by Eq. (S42).
